# Tvp complexes support formation of the VgrG-PAAR spike during Type VI secretion system assembly

**DOI:** 10.64898/2026.02.23.707444

**Authors:** Laura Monlezun, Christopher Earl, Sarah J. Coulthurst

**Affiliations:** School of Life Sciences, University of Dundee, Dow Street, Dundee DD1 5EH, UK; Université Paris Cité, CNRS, Expression Génétique Microbienne, Institut de Biologie Physico-Chimique, Paris, France

## Abstract

Type VI secretion systems (T6SSs) are nanomachineries used by bacteria to inject toxic effectors into neighbouring cells during interbacterial competition or infection. The existence of multiple paralogues of the structural proteins VgrG and PAAR and of specific accessory proteins allows the same T6SS to deliver a wide range of effectors. We describe a new set of accessory proteins required for assembly of T6SSs containing the VgrG1-Paar1 spike and delivery of a novel VgrG1-associated membrane-targeting effector in *Serratia marcescens*. TvpAB, TvpB, TvpC and Paar1 form a pre-complex essential for T6SS assembly, with VgrG1 subsequently replacing TvpC. Phylogenetic analysis and structural modelling reveal that Tvp proteins are widely conserved but Tvp-containing complexes vary in organisation and complexity. We define three classes of Tvp complex, all sharing a DUF2169-containing TvpA protein which stabilises a DUF4150/PAAR-containing protein. Class I, which includes only TvpA, functions without a pre-complex, whereas Classes II and III involve additional Tvp components and two-step assembly. Our findings highlight how modular effector recruitment strategies underlie the versatility of the T6SS and suggest how alternative Tvp systems are tailored to promote assembly of secretion-competent, effector-loaded VgrG-PAAR spikes.

## INTRODUCTION

The Type VI secretion system (T6SS) is a contractile nanomachine used by many Gram-negative bacteria to deliver toxic effectors into neighbouring cells. Although it can mediate intoxication of eukaryotic cells or promote the secretion of metal-scavenging proteins, its major role lies in intra- and inter-species competition between bacteria^1^. The T6SS apparatus comprises 14 core components in four sub-complexes: the baseplate, the membrane complex, the sheath, and an expelled puncturing structure. T6SS assembly starts with the anchoring of the membrane complex (TssJ, TssL, TssM) into the cell envelope, followed by the recruitment of the baseplate (VgrG/PAAR, TssE, TssF, TssG, TssK) to the inner face of the cytoplasmic membrane. The trimeric VgrG spike protein, sharpened by a conical PAAR protein tip, forms the central hub of the baseplate and its base serves as the scaffold for the polymerisation of a tube of stacked hexameric Hcp rings. Extending into the cytoplasm, this spear-like puncturing structure, comprising the VgrG-PAAR spike topping the Hcp tube, is surrounded by the sheath, composed of TssB and TssC subunits. Upon contraction of the sheath, also known as ‘firing’, the puncturing structure, loaded with effector proteins, is propelled across the envelope of the secreting bacterium and perforates the target cell to deliver the effectors^2^.

T6SS effectors can be categorised based on their mode of delivery. Cargo effectors interact non-covalently with VgrG, PAAR or Hcp proteins within the puncturing structure, whilst specialised effectors comprise effector domains covalently attached to the C-terminus of one of these proteins^3^. Certain VgrG proteins contain C-terminal extensions that are not toxins but are required for the recruitment of other effectors. VgrG from EAEC, for example, includes a DUF2345 domain and a TTR domain which are both important for Tle1 delivery^4^. Additionally, adaptor proteins can be required for the loading of T6SS effectors onto the machinery. Two well-characterised classes of adaptor have been described: DUF4123-family proteins, which interact with a VgrG C-terminal extension and their cognate cargo effector to facilitate loading^5^, and DUF1795-containing (‘Eag’) chaperone proteins which load PAAR-containing specialised effectors onto their cognate VgrG protein whilst protecting their transmembrane domains^6^. Several studies have indicated that DUF2169-containing proteins may also function as T6SS adaptor proteins. In *Agrobacterium tumefaciens*, Atu3641/Tap2 was shown to be required for VgrG2-dependent delivery of the PAAR-containing Tde2 effector^7^, with a subsequent study reporting that genetic loci containing *vgrG and tap2* genes always encode a protein containing a DUF4150 (PAAR-like) domain and usually also an effector domain^8^. Recently, an experimental structure was determined for a DUF2169 protein, and it was shown to interact with a DUF4150 protein to facilitate the formation of a functional VgrG-PAAR complex in *Vibrio*^10^.

*Serratia marcescens* is an opportunistic bacterium found in various environmental niches and also responsible for causing hospital-acquired infections^11^. The model *S. marcescens* strain Db10 possesses a single T6SS which displays anti-bacterial and anti-fungal activities^12,13^. Ten T6SS effectors have been identified so far in Db10, three encoded in the main T6SS gene cluster and the others elsewhere in the genome^13^. The *S. marcescens* T6SS is associated with two VgrG proteins, VgrG1, encoded at the 5’ end of the T6SS gene cluster, and VgrG2 at the 3’ end, and can form three distinct assemblies based on specific VgrG-PAAR combinations^14^. VgrG1 functions with the DUF4150-type Paar1 protein, whereas VgrG2 can function with two different specialised PAAR effectors, Rhs1 or Rhs2. Both the VgrG1- and VgrG2-containing T6SS assemblies can deliver Hcp-dependent cargo effectors (Ssp1-Ssp6, Tfe2), however, VgrG2-based assemblies deliver cargo effectors more efficiently than VgrG1-Paar1. Interestingly, whilst another three effectors, Rhs1, Rhs2, and Slp, are exclusively delivered by VgrG2, VgrG1-specific effectors remain to be identified^14^.

The 5’ end of the *S. marcescens* Db10 T6SS gene cluster contains five genes of unknown function (*SMDB11_2245-2249*) located between *vgrG1* and *paar1* (**Figure 1a**), suggesting that they could be involved in delivery of T6SS effectors by the VgrG1-Paar1 assembly. Here, we show that SMDB11_2247, SMDB11_2248, SMDB11_2249 and Paar1 form an accessory complex required for the VgrG1 pathway, which delivers a novel membrane-targeting anti-bacterial effector against which SMDB11_2245 provides immunity. Based on their newly identified functions, we name SMDB11_2247-2249 as TvpAB, TvpB and TvpC, for <u>T</u>ag protein chaperoning <u>V</u>grG-<u>P</u>AAR complexes. Through phylogenetic analysis and structural modelling, we define three classes of Tvp system, all of which are proposed to have the common function of assembling a mature VgrG-PAAR spike. The Class I system is the simplest, represented by the VgrG-TvpA-PAAR complex of *A. tumefaciens*. Classes II and III require an intermediary TvpC-containing complex, which templates the formation of the final VgrG-containing accessory complex. Class II systems, represented by the VgrG1b-associated complex from *Pseudomonas aeruginosa*, are characterized by a specific family of TvpC proteins (DUF6484) and the presence of an additional component named TvpD. Class III, the most complex, is represented by the *S. marcescens* system studied experimentally here, with longer TvpC proteins (DUF3540) and pentapeptide repeat-containing TvpB proteins. Our findings reveal that assembly of a secretion-competent, effector-loaded T6SS spike and resulting effector delivery is more complex than previously appreciated and can involve multiple steps. We propose that Tvp accessory complexes both support interactions between specific VgrG and PAAR proteins and fulfil a requirement to sufficiently occupy the cavity around the spike in the baseplate.

**Figure 1.**
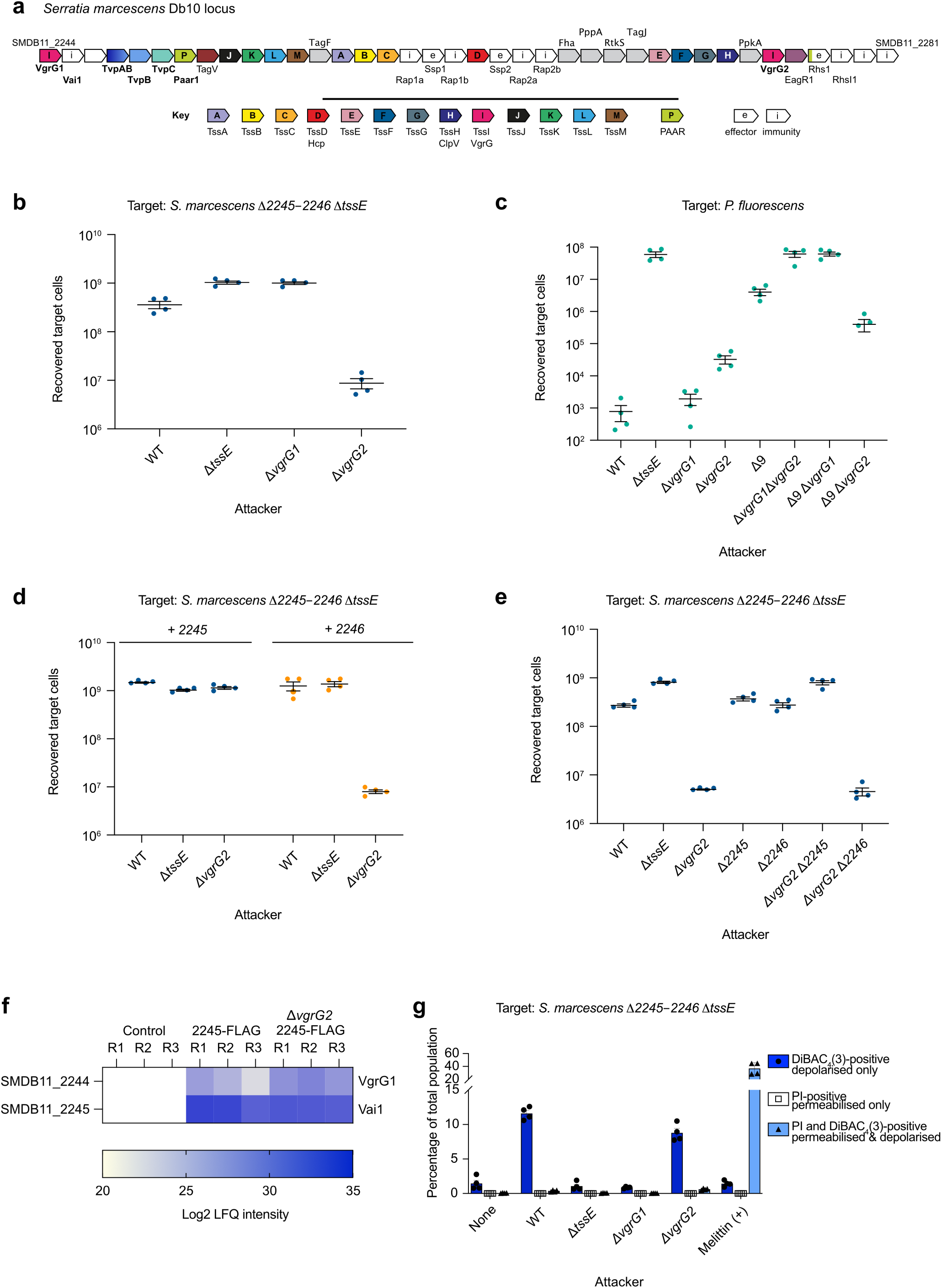
VgrG1 is responsible for a unique anti-bacterial activity that causes depolarisation of the target cell membrane and against which Vai1 provides immunity. **a)** Schematic of the main T6SS gene cluster in *S. marcescens* Db10. Genes encoding core T6SS components are coloured and labelled with their corresponding Tss letters. **b,e)** Recovery of *S. marcescens* Db10 Δ*2245-2246* Δ*tssE* target cells following co-culture with wild type (WT) and mutant strains of *S. marcescens* Db10 at an initial ratio of 1:1. **c)** Recovery of *P. fluorescens* target cells following co-culture with WT and mutant strains of *S. marcescens* at an initial ratio of 1:1. **d)** Recovery of the Δ*2245-2246* Δ*tssE* target complemented with either *SMDB11*_*2245* or *SMDB11*_*2246* expressed from the pSUPROM plasmid, following co-culture with WT and mutant strains of *S. marcescens* at an initial ratio of 1:1. **f)** Heatmap of label-free proteomic quantification (LFQ) intensity values for proteins in the elution fractions of anti-FLAG immunoprecipitations from the control strain (WT Db10, untagged) or from strains expressing an N-terminal 3xFLAG fusion to SMDB11_2245 (Vai1) from the normal chromosomal location, in either the parental background (FLAG-2245) or Δ*vgrG2* (ΔvgrG2 FLAG-2245). Only proteins exclusively detected or significantly enriched (log2 fold change >2, p<0.05) in immunoprecipitation samples compared with the control samples are included. The experiment included three biological replicates of each strain (R1, R2, R3). **g)** The VgrG1-susceptible target (Δ*2245-2246* Δ*tssE*) was co-cultured with WT and mutant strains of *S. marcescens* and then membrane potential and membrane permeability of the mixed population was determined by staining with DiBAC_4_(3) and propidium iodide (PI) followed by flow cytometry analysis. The percentage of cells in the total co-culture population identified as being permeabilised only (positive for fluorescence from PI only), depolarised only (positive for fluorescence from DiBAC_4_(3) only), or simultaneously depolarised and permeabilised (positive for PI and DiBAC_4_(3)) is shown on the Y-axis. Bars show mean +/− SEM, with individual data points overlaid (n=4 independent experiments). **b-e)** Data sets are displayed as mean +/− SEM (n=4) with individual data points overlaid.

## RESULTS

### A novel VgrG1-dependent effector compromises the membrane potential of intoxicated bacterial cells and is neutralised by the immunity protein Vai1

Since no T6SS effectors specific to VgrG1 had been identified in *S. marcescens* Db10, we considered whether two proteins of unknown function encoded directly downstream of *vgrG1*, SMDB11_2245, which contains a DUF3192 domain, and SMDB11_2246, represented a new effector-immunity pair (**Figure 1a**). We generated a Δ*2245-2246* Δt*ssE* mutant and assessed its susceptibility to T6SS-mediated anti-bacterial activity. Co-culture of this target with wild type Db10 resulted in reduced recovery of viable target cells compared to co-culture with a T6SS mutant (Δ*tssE*) (**Figure 1b**), indicating that the Δ*2245-2246* mutant lacks immunity against a T6SS-delivered effector. Deletion of *vgrG1* in the attacker cells abolished inhibition of the target, while deletion of *vgrG2* greatly increased the anti-bacterial activity towards this target, consistent with our previous findings that secretion of VgrG1 is increased in the absence of VgrG2^14^. Moreover, the Δ9 Δ*vgrG2* mutant, lacking all known antibacterial effectors in Db10, was indistinguishable from Δ*vgrG2*, confirming that known effectors are not responsible for activity against Δ*2245-2246* (**Supplementary Figure 1a**). Using *Pseudomonas fluorescens* as a target, we next showed that the remaining antibacterial activity of the Δ9 strain, compared with a T6SS mutant, was dependent on VgrG1, being abolished in a Δ9 Δ*vgrG1* mutant. Recovery of *P. fluorescens* was 2-log lower using the Δ9 Δ*vgrG2* attacker strain compared with the T6SS mutant, also showing that the VgrG1-dependent effector can have considerable efficacy in an interspecies competition setting (**Figure 1c**). Overall, these results show that *P. fluorescens* and the *S. marcescens* Δ*2245-2246* mutant are susceptible to a previously-unidentified VgrG1-dependent effector and that the delivery of this effector is increased in the absence of VgrG2, *i.e.,* when there is no competition between the two VgrGs for assembly onto the T6SS machinery.

In order to determine which gene encodes the immunity protein, the intra-species co-culture experiment was repeated using a target with *SMDB11_2245* or *SMDB11_2246* deleted individually in a Δ*tssE* background. The Δ*2245* Δ*tssE* mutant remained sensitive to the antibacterial activity of the Δ*vgrG2* strain, whereas the Δ*2246* Δ*tssE* mutant was resistant (**Supplementary Figure 1b,c**), indicating that SMDB11_2245 is the immunity protein conferring resistance to the effector delivered by the VgrG1 pathway. In addition, *in trans* complementation of the Δ*2245-2246* Δt*ssE* target strain confirmed that only SMDB11_2245 could provide protection against killing by the Δ*vgrG2* attacker strain (**Figure 1d**). In an attempt to identify the cognate effector, Δ*2245* and Δ*2246* single mutants were generated in a wild type or Δ*vgrG2* background and tested as attackers against the Δ*2245-2246* Δ*tssE* target strain. Deletion of *SMDB11_2246* had no impact on activity against the Δ*2245*-Δ*2246* Δ*tssE* target (**Figure 1e**), indicating that SMDB11_2246 is not the cognate effector, despite being encoded next to the immunity gene. Deletion of *SMDB11_2245* led to a reduction in antibacterial activity in the wild type and, particularly, the Δ*vgrG2* background, likely due to self-toxicity in the absence of the immunity protein.

In order to identify the effector to which SMDB11_2245 provides immunity, a co-immunoprecipitation experiment was carried out using SMDB11_2245 with a C-terminal 3xFLAG tag (2245-FLAG) as bait, having first verified that the 3xFLAG tag did not impair the protective function of SMDB11_2245 (**Supplementary Figure 1d**). The immunoprecipitation was conducted under the same conditions as the antibacterial co-culture assays and in a wild type and Δ*vgrG2* background. The only protein identified in the immunoprecipitation, besides 2245-FLAG, was VgrG1 (**Figure 1f**). This suggested that the effector function may reside within VgrG1, i.e. that VgrG1 might be a specialised effector with an effector domain at its C-terminus. VgrG1 contains 786 amino acids, 144 more than VgrG2, thus this extension could represent an effector domain. However, no recognisable effector domain is present in this region (see below) and multiple attempts to isolate the toxic activity of VgrG1 in a heterologous expression system proved unsuccessful. Therefore, it is possible that the firing of an intact VgrG1 into a target cell by the T6SS is vital for the mechanism of toxicity.

To gain insight into the nature of the VgrG1-dependent effector, TMHMM and AlphaFold predictions for the immunity protein were generated in order to identify the cellular compartment in which the effector acts. SMDB11_2245 was predicted to have an N-terminal transmembrane helix (aa 7-29) in the inner membrane, with the body of the protein located in the periplasm and primarily composed of β-strands (**Supplementary Figure 1e**). This localisation, together with the observation that the Δ*2245-2246* Δ*tssE* mutant does not intoxicate itself, suggests that the cognate toxin either acts within the periplasm or on the inner membrane from the periplasm. In order to investigate the impact of VgrG1-dependent toxicity on the inner membrane of intoxicated bacterial cells, we co-cultured attacker cells delivering the effector with target cells susceptible to the effector (Δ*2245-2246*), then stained the cells with propidium iodide (PI) and DiBAC_4_(3) and analysed the total population by flow cytometry. DiBAC_4_(3) stains cells which have lost their inner membrane potential (depolarised cells), for example if ions can flow freely across the membrane, whilst PI staining occurs upon formation of large, non-specific pores or loss of membrane integrity (permeabilised cells). Target cells were also treated with melittin, a natural anti-bacterial peptide forming pores in cell membranes^15^, as a control (**Figure 1g and Supplementary Figure 2a**). Use of Db10 or Δ*vgrG2* as attackers induced membrane depolarisation, with around 10% of the total population stained with DiBAC_4_(3) only, but this effect was not observed with the Δ*tssE* and Δ*vgrG1* attackers (**Figure 1g and Supplementary Figure 2b**). The absence of an increase in DiBAC_4_(3)-positive cells using Δ*vgrG2* as the attacker compared with Db10 can be explained by a significant reduction in the number of recovered target cells in this condition, even with a reduced attacker:target ratio and incubation time compared with the standard co-culture setting (**Supplementary Figure 2c**). Altogether, these data reveal that the VgrG1 pathway delivers a further antibacterial activity, either within or interacting with VgrG1, which distinguishes it from the VgrG2 pathway. Toxicity is mediated via an effect on the inner membrane of target cells, likely the formation of small ion channels. Based on the newly identified immunity function of SMDB11_2245 against this VgrG1-specific anti-bacterial effector, we name SMDB11_2245 Vai1 (<u>V</u>grG1-<u>a</u>ssociated immunity protein).

### TvpAB, TvpB and TvpC accessory proteins are essential for a functional VgrG1-Paar1 assembly

The presence of genes encoding three other proteins, SMDB11_2247, SMDB11_2248 and SMDB11_2249, between *vgrG1* and *paar1* in *S. marcescens* Db10 (**Figure 1a**) prompted us to investigate their possible role in the VgrG1 delivery pathway. Single in-frame deletion mutants of each gene were tested in a co-culture assay against the VgrG1-susceptible target, Δ*2245*-Δ*2246* Δ*tssE*. Loss of SMDB11_2247, SMDB11_2248, SMDB11_2249 or Paar1 abolished activity against this target strain in the same manner as deletion of *vgrG1* or *tssE* (**Figure 2a**). In an interspecies competition, recovery of *P. fluorescens* was not affected by deletion of *SMDB11_2247*, *SMDB11_2248, SMDB11_2249* or *paar1* genes in a wild type background, since the more potent VgrG2 delivery pathway is still functional. However, antibacterial activity was fully abolished for these mutants in the absence of VgrG2, *i.e.*, when only the VgrG1 pathway is present (**Figure 2b**). The requirement of these genes for the functionality of the VgrG1 pathway was verified by examining Hcp secretion, a direct proxy of T6SS firing. Consistent with the existence of distinct functional T6SS assemblies in *S. marcescens* depending on the VgrG-PAAR pairing, neither the individual deletion of *vgrG1* or *vgrG2* abolished secretion of Hcp, although secreted Hcp levels were slightly lower in the Δ*vgrG2* strain and absent, as expected, in the Δ*vgrG1*Δ*vgrG2* mutant (**Figure 2c**). In agreement with the results observed for T6SS-dependent antibacterial activity (**Figure 2b**), deletion of *SMDB11_2247, SMDB11_2248, SMDB11_2249* or *paar1* genes prevented Hcp secretion in the absence of VgrG2 (**Figure 2c**). Finally, complementation of these mutants by expression of the respective genes *in trans* restored T6SS activity against the *S. marcescens* Δ*2245-2246* Δ*tssE* target (**Figure 2d**). Based on their essential accessory role in the VgrG1-Paar1 pathway, and an early study naming homologous genes *tagAB*, *tagB* and *tagC* (*tag*, tss-associated gene)^49^, SMDB11_2247, SMDB11_2248 and SMDB11_2249 were renamed TvpAB, TvpB and TvpC, respectively, for <u>T</u>ag protein chaperoning <u>V</u>grG-<u>P</u>AAR complex.

**Figure 2.**
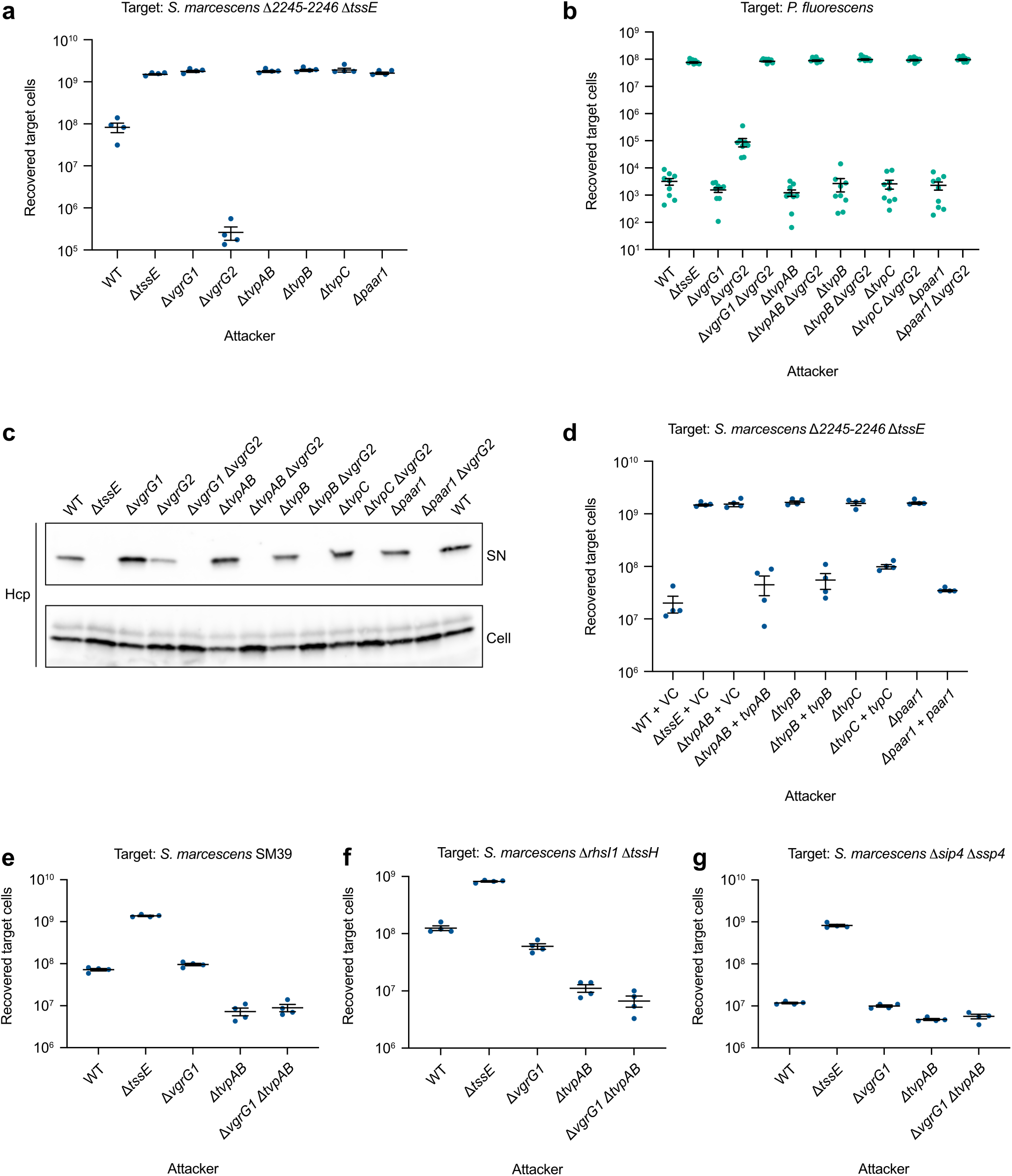
TvpAB, TvpB and TvpC are accessory proteins required for the VgrG1-Paar1 delivery pathway. **a)** Recovery of *S. marcescens* Db10 Δ*2245-2246* Δ*tssE* target cells following co-culture with wild type (WT) and mutant strains of *S. marcescens* Db10 at an initial ratio of 1:1. **b)** Recovery of *P. fluorescens* target cells following co-culture with wild type and mutant strains of *S. marcescens* at an initial ratio of 1:1. **c)** Levels of Hcp1 in total cellular (Cell) and secreted (SN) protein fractions from the indicated strains of *S. marcescens* as detected by immunoblot. **d)** Recovery of *S. marcescens* Δ*2245-2246* Δ*tssE* target cells following co-culture with WT and mutant strains of *S. marcescens* carrying either the vector control plasmid (pSUPROM, VC) or plasmids directing the expression of TvpAB, TvpB, TvpC or PAAR *in trans*, at an initial ratio of 1:1. **e-g)** Recovery of *S. marcescens* SM39 (**e**), Db10 Δ*rhsI1* Δ*tssH* (**f**), or Db10 Δ*sip4* Δ*ssp4* (**g**), target cells following co-culture with WT and mutant strains of *S. marcescens* Db10 at an initial ratio of 1:1. Data sets are displayed as mean +/− SEM (n=4) with individual data points overlaid.

Initial co-culture assays against *P. fluorescens* suggested there might be a slight increase in activity with the Δ*tvpAB* attacker strain compared with the wild type (e.g. **Supplementary Figure 3**). Thus, we tested other targets, namely another strain of *S. marcescens* and two derivatives of Db10 sensitive to the Rhs1 and Ssp4 effectors, and observed that the Δ*tvpAB* mutant showed a clear increase in activity against these targets (**Figure 2e-g)**. Interestingly, such an increase was not observed with the Δ*vgrG1* mutant. The fact that only the Δ*tvpAB* mutant, but not the Δ*vgrG1* mutant, presents this phenotype indicates that the effect is not simply due to competition for the T6SS machinery between the VgrG1 pathway and the more efficient VgrG2 pathway. Moreover, enhanced activity was also observed for the Δ*tvpAB* Δ*vgrG1* mutant (**Figure 2e-g and Supplementary Figure 3**) confirming that the TvpAB repressive effect directly affects the VgrG2 pathway and is independent of the VgrG1 pathway. However, no protein-protein interactions between TvpAB and VgrG2 pathway components have been detected to explain this additional function of TvpAB^14^ (and below). Nevertheless, these findings strongly suggest that TvpAB mediates interference between the VgrG1 and VgrG2 pathways, although the underlying mechanism remains to be elucidated.

### The VgrG1 accessory complex requires the formation of a TvpC pre-complex

The requirement for TvpAB, TvpB and TvpC in the VgrG1 delivery pathway suggested a potential interaction between those proteins and the T6SS spike. A strain encoding VgrG1 fused with an N-terminal His_6_-tag at the native chromosomal location was constructed and functionality of the fusion protein confirmed (**Supplementary Figure 4a**). His-VgrG1 was then affinity-purified from cell lysates and co-purifying proteins identified by quantitative mass spectrometry. Only three proteins were significantly enriched in the His-VgrG1 pulldown compared to the control, namely TvpAB, TvpB, and Paar1 (**Figure 3a**). Interestingly, although TvpC was essential for the functionality of the VgrG1 pathway (**Figure 2a-d**), this protein was not detected among the VgrG1 interacting partners (**Figure 3a**). To understand this discrepancy, we generated a strain encoding an N-terminally HA-tagged version of TvpC and confirmed its functionality (**Supplementary Figure 4b**). Identification of proteins significantly enriched in an HA-TvpC co-immunoprecipitation revealed TvpAB, TvpB, and Paar1 as the strongest hits (**Figure 3b**). Once again, no interaction between TvpC and VgrG1 was detected. These results suggest the existence of two different complexes, a VgrG1 accessory complex and a TvpC alternative complex, which are mutually exclusive but share the same components, namely TvpAB, TvpB and Paar1. Consistent with this finding, when TvpAB with an HA tag at the C-terminus (**Supplementary Figure 4c**) was used as bait (**Figure 3c**), all the components (Paar1, TvpB, TvpC and VgrG1) were co-purified. Importantly, deletion of either TvpAB or TvpC prevented the formation of the VgrG1 accessory complex (**Figure 3a**), while deletion of VgrG1 did not impact the formation of the alternative TvpC complex, since TvpAB, TvpB, TvpC, and Paar1 were still co-purified in Δ*vgrG1* TvpAB-HA and Δ*vgrG1* HA-TvpC immunoprecipitations (**Figure 3b-c**). This indicates that formation of the TvpC alternative complex is a prerequisite for the formation of the VgrG1 accessory complex. Moreover, deletion of TvpAB also abolished the formation of the TvpC alternative complex (**Figure 3b**), while deletion of TvpC considerably reduced the amount of bait recovered from the TvpAB-HA immunoprecipitation (**Figure 3c**). Overall, our interaction studies identify a TvpC-containing pre-complex (TvpC-TvpAB-TvpB-Paar1) which is required for the subsequent formation of the VgrG1-containing accessory complex (VgrG1-TvpAB-TvpB-Paar1), with all of these components being essential for assembly and firing of T6SS machineries containing VgrG1-Paar1 as the spike complex.

**Figure 3.**
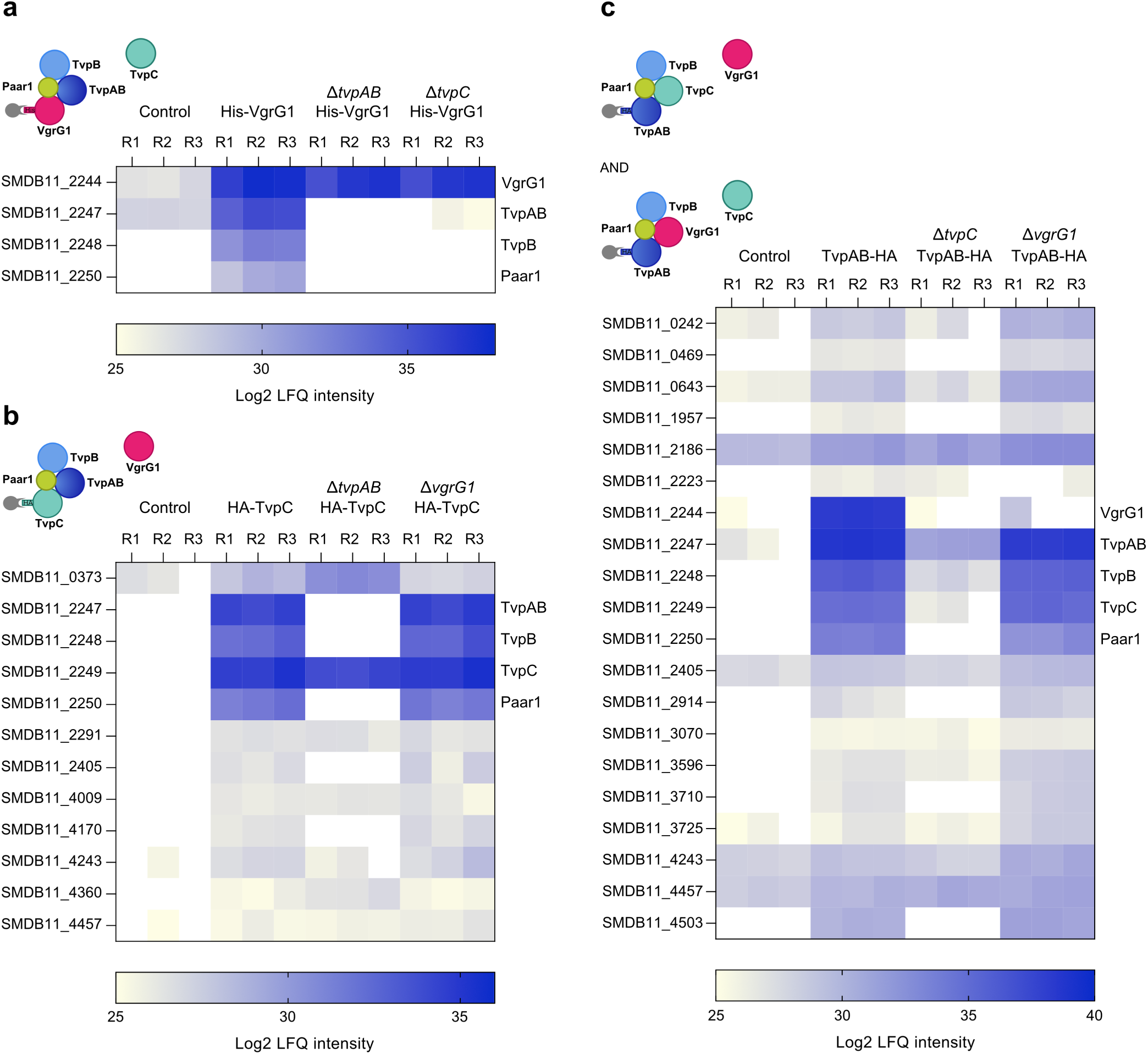
TvpAB, TvpB and Paar1 interact with TvpC and with VgrG1 but the presence of TvpC and VgrG1 in these complexes is mutually exclusive. **a)** Heatmap of label-free proteomic quantification (LFQ) intensity values for proteins in the elution fractions of Ni-affinity purifications from the control strain (WT Db10, untagged) or from strains expressing a His_6_-VgrG1 fusion protein from the normal chromosomal location, in the parental (His_6_-VgrG1), Δ*tvpAB* (Δ*tvpAB* His_6_-VgrG1) or Δ*tvpC* (Δ*tvpC* His_6_-VgrG1) genetic backgrounds. **b)** Heatmap of LFQ intensity values for proteins in the elution fractions of anti-HA immunoprecipitations from the control strain or from strains expressing an HA-TvpC fusion protein from the normal chromosomal location, in the parental (HA-TvpC), Δ*tvpC* (Δ*tvpC* HA-TvpC) or Δ*vgrG1* (Δ*vgrG1* HA-TvpC) genetic backgrounds. **c)** Heatmap of LFQ intensity values for proteins in the elution fractions of anti-HA immunoprecipitations from the control strain or from strains expressing a TvpAB-HA fusion protein from the normal chromosomal location, in the parental (TvpAB-HA), Δ*tvpC* (Δ*tvpC* TvpAB-HA) or Δ*vgrG1* (Δ*vgrG1* TvpAB-HA) genetic backgrounds. In each case, only proteins exclusively detected or significantly enriched (log2 fold change >2, p<0.05) in affinity purification or immunoprecipitation samples compared with the control samples are included in the heat maps. Each experiment included three biological replicates of each strain (R1, R2, R3). A schematic of the complexes identified in each pulldown/IP is depicted on the top left corner of the heatmaps. Full mass spectrometry data for each experiment are included in Supplementary Dataset 1.

### TvpC shares structural homology with VgrG proteins

To understand why the VgrG1 accessory complex and TvpC alternative complex are mutually exclusive while sharing the same partners, a structural prediction of TvpC was generated using AlphaFold2 and compared with predictions of VgrG1 and VgrG2. The TvpC protein is predicted to form a trimer and is composed of two domains, both folded with a high confidence score (pLDDT >90) (**Figure 4**). These domains, an OB-fold and a β-prism, are also found in VgrG proteins. The main difference compared with VgrG proteins is the absence, in TvpC, of the Gp27 domain which interacts with the Hcp tube, and its replacement by a 16 amino acid unstructured tail at the N-terminus. This further suggests that the TvpC complex is an intermediate during the assembly of the T6SS machinery. Another striking feature of the structural comparison between the three proteins is the length of the β-prism. While TvpC and VgrG2 both have a β-prism 81 Å in length, VgrG1 has a very long β-prism of 186 Å (**Figure 4**). Although the confidence score is somewhat lower for the second half of this long β-prism (pLDDT >70), it appears that the entire C-terminus of VgrG1 is part of this domain. Therefore, the VgrG1 C-terminus does not form a specific effector domain as is found in classical specialised VgrGs and which could readily explain the VgrG1-associated toxicity characterised above. Moreover, an AlphaFold prediction of the VgrG1-Paar1 complex confirms the presence of the long β-prism, whose tip seems to be further stabilized by the Paar1 protein, as evidenced by a higher pLDDT score for the VgrG1 extremity when complexed with Paar1 than when alone (**Supplementary Figure 5**).

**Figure 4.**
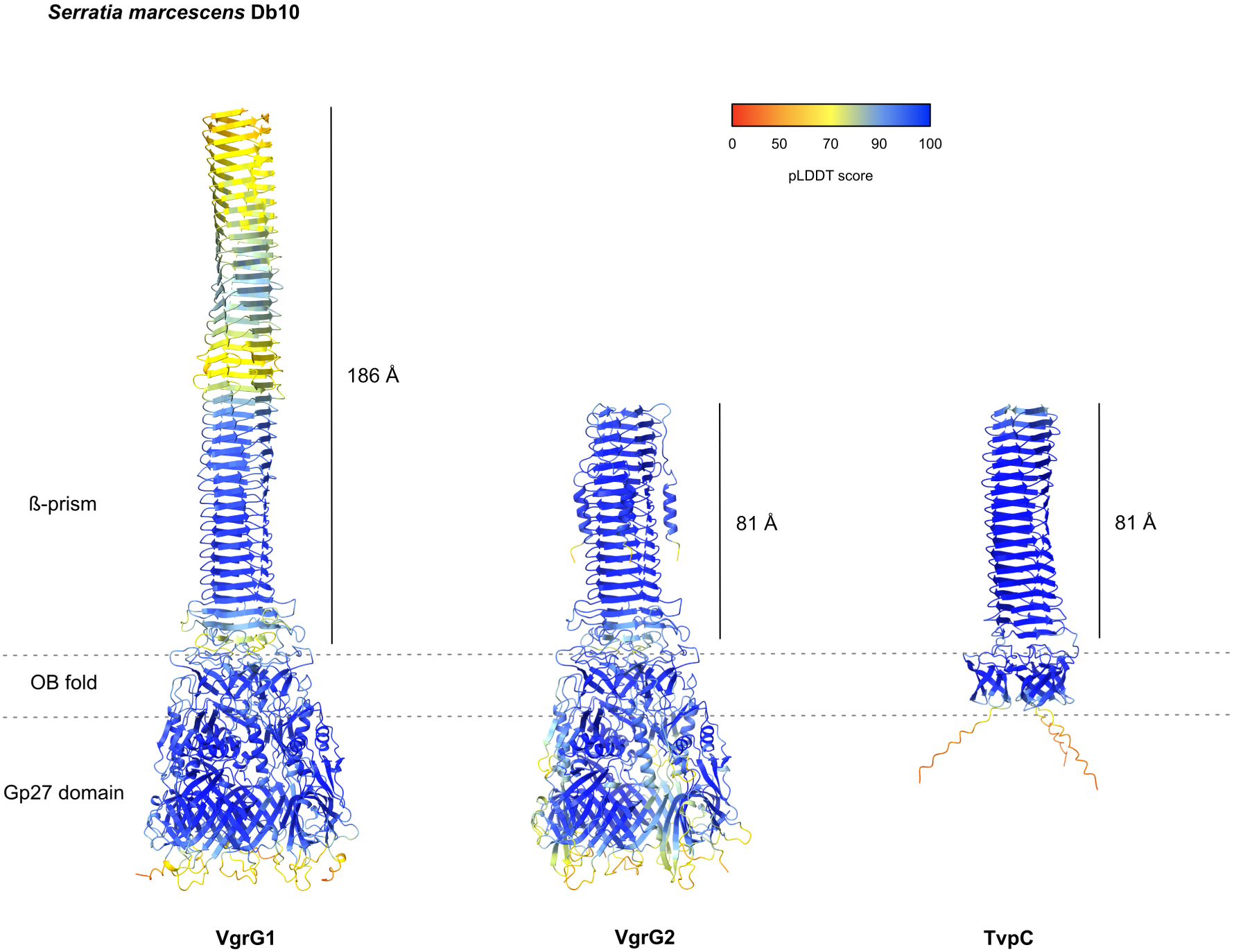
TvpC shares structural homology with VgrG proteins. Predicted structures of VgrG1, VgrG2 and TvpC from *S. marcescens* Db10. Ribbon representations are coloured according to the reported confidence of the AlphaFold2 modelling, from red (pLDDT <50, lowest confidence) to blue (pLDDT >90, highest confidence). The length of the β-prism is indicated next to each structure prediction.

### Tvp accessory proteins are conserved in other species with a functional T6SS

The remarkable length of the VgrG1 β-prism prompted us to investigate whether such a characteristic could explain the requirement for the Tvp accessory proteins. We looked for VgrG-associated accessory genes in other bacterial genomes, focusing on bacteria with an experimentally-validated T6SS and whose genomes contain a gene encoding a protein with a DUF2169 domain. We discovered that a number of other organisms, including *Pantoea ananatis*, *Cronobacter sakazakii*, and *Burkholderia thailandensis* or *pseudomallei,* possess the same genetic organization of *tvp-paar* accessory genes downstream of their *vgrG* genes as *S. marcescens* (**Figure 5a** and **Supplementary Figure 6**). In addition, other organisms, like *A. tumefaciens*, harbour only a gene encoding a protein with a DUF2169 domain followed by a specialised PAAR effector gene, while others, like *P. aeruginosa* or *Proteus mirabilis*, possess two other putative accessory genes: a gene encoding a protein with a DUF6484 domain located downstream of *vgrG* and a gene encoding a protein with a PRK06147 domain located between the DUF2169 gene and a PAAR-containing effector gene (**Figure 5a** and **Supplementary Figure 6**). In these two alternative configurations, the DUF2169-containing gene encodes a shorter homologue of TvpAB which lacks the PPR domain and therefore can be described as a TvpA protein, whilst no homologue of TvpB could be identified. In all of these *tvp*-like clusters, the *vgrG* genes encode VgrG proteins which are one of at least two VgrGs associated with that T6SS and which are usually 700-800 amino acids long, except for VgrG5 from *Burkholderia* which has a characterised C-terminal effector domain^16^. Therefore, the presence of Tvp proteins seems to be associated with non-essential and rather long VgrG proteins, and their role in all these organisms is likely to be to chaperone alternative VgrG-PAAR assemblies.

**Figure 5.**
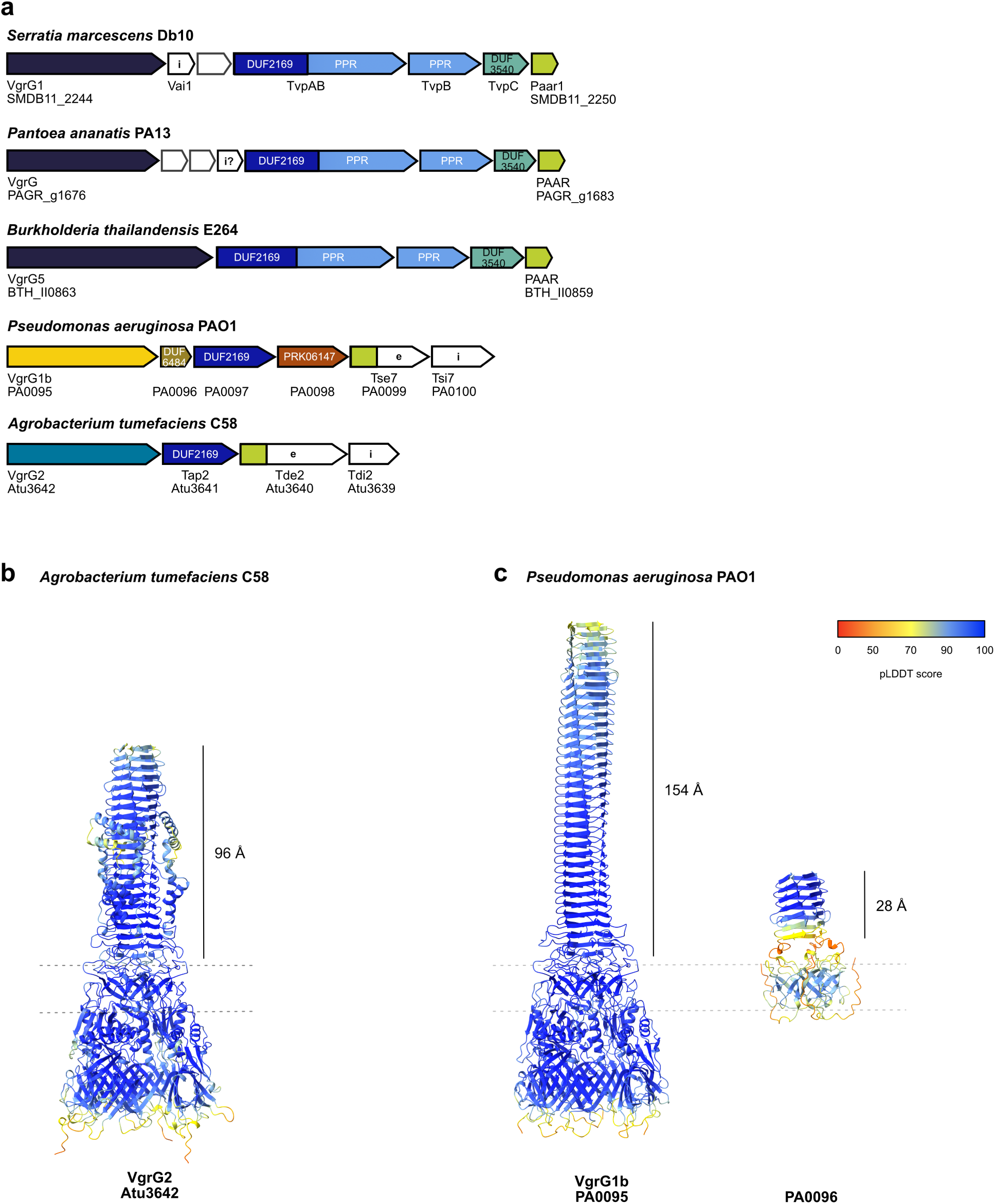
Tvp accessory proteins are conserved in many organisms with a functional T6SS. **a)** Genetic context of Tvp and other VgrG accessory proteins in *Pantoea ananatis* PA13 (accession: CP003085.1), *Burkholderia thailandensis* E264 (accession: CP000085.1 and CP000086.1), *Pseudomonas aeruginosa* PAO1 (accession: AE004091.2) and *Agrobacterium tumefaciens* C58, now known as *A. fabrum* C58 (accession: AE007869.2). A larger view of the conservation of Tvp accessory proteins in other organisms is provided in Supplementary Figure 6. **b)** Predicted structure of VgrG2 from *A. tumefaciens* C58. **c)** Predicted structures of VgrG1b and PA0096 from *P. aeruginosa* PAO1. **b-c)** Predicted structures were generated by AlphaFold2 and ribbon representations are colored according to pLDDT score, from red (pLDDT <50, lowest confidence) to blue (pLDDT >90, highest confidence). The length of the β-prism is indicated next to each structure prediction.

Based on the existence of these three distinct organizations of *tvp* accessory genes associated with *vgrG*, we used AlphaFold2 to predict the structure of the *tvp*-associated VgrG proteins from *A. tumefaciens* and *P. aeruginosa* as well as the structures of their respective Tvp-like accessory proteins. VgrG2 from *A. tumefaciens* (Atu3642) has a 96 Å β-prism (**Figure 5b**) while VgrG1b from the H1-T6SS of *P. aeruginosa* (PA0095) has a 154 Å β-prism (**Figure 5c left**), making them intermediate in their length between VgrG1 and VgrG2 from *S. marcescens*. Both of their TvpA-like accessory proteins, namely Tap2 for *A. tumefaciens* (Atu3641) and PA0097 for *P. aeruginosa*, are predicted to adopt the same fold as the DUF2169 domain of TvpAB (**Supplementary Figure 7b-d**). Strikingly, PA0096 from *P. aeruginosa*, which contains the DUF6484 domain, adopts a fold similar to TvpC of *S. marcescens* although its β-prism is only 28 Å long (**Figure 5c right**). This implies that the Tvp-like accessory proteins from *P. aeruginosa* likely also form an alternative pre-complex before interacting with VgrG1b.

### Predicted structures of accessory pre-complexes and VgrG-associated full complexes

To gain insight into how the Tvp accessory proteins may chaperone VgrG1-Paar1 assembly, we first used AlphaFold2 to predict the structure of the TvpC pre-complex. This pre-complex was successfully modelled with a high confidence score (pLDDT >90) (**Supplementary Figure 8a**). The predicted structure shows that Paar1 interacts with the extremity of the TvpC β-prism (**Figure 6a**), similar to its interaction with VgrG. TvpAB then wraps around the TvpC-Paar1 into its concave groove, while TvpB acts as a “seat belt” to fasten the pre-complex (**Figure 6a and Supplementary Figure 8b**). Thus, the tip of the TvpC-Paar1 structure is completely enfolded by the TvpAB-TvpB tandem. In this pre-complex, the DUF2169 domain of TvpAB interacts with Paar1, with surface analysis indicating that the interaction is mainly driven by hydrophobic regions of the two partners (**Supplementary Figure 9a**). Other hydrophobic patches of Paar1 are masked by hydrophobic parts of the TvpB PPR domain. As a comparison, we also obtained an AlphaFold prediction of the *P. aeruginosa* pre-complex formed by PA0096 (DUF6484, TvpC-like), PA0097 (TvpA), PA0098 (PRK06147) and the PAAR domain of Tse7. The confidence score of this pre-complex is slightly lower compared to *S. marcescens* (pLDDT >80) but the uncertainty is mainly concentrated on the extremities of PA0096 and PA0097 (**Supplementary Figure 8c**). In this predicted structure, the PAAR domain of Tse7 sits on the top of PA0096 β-prism, as in *S. marcescens*, while PA0097 and PA0098 form a concave groove protecting the PAAR domain of Tse7 (**Figure 6b** and **Supplementary Figure 8d)**. However, this chaperoning tandem is more open than in *S. marcescens*. The PA0097 DUF2169 domain shields the hydrophobic surface of the PAAR domain of Tse7 while the Tse7 hydrophilic face remains unprotected (**Supplementary Figure 9b**). A structural prediction with the full sequence of Tse7 indicates that the effector domain does not close the belt around the PAAR domain but rather protrudes away from the pre-complex, outside of the PA0097-PA0098 face (**Supplementary Figure 10**). In addition, in the *P. aeruginosa* pre-complex, PA0096, the OB-fold/TvpC protein, barely interacts with PA0097 containing the DUF2169 domain or with PA0098, since both of these proteins are shorter than their homologues in *S. marcescens* and devoid of a PPR domain.

**Figure 6.**
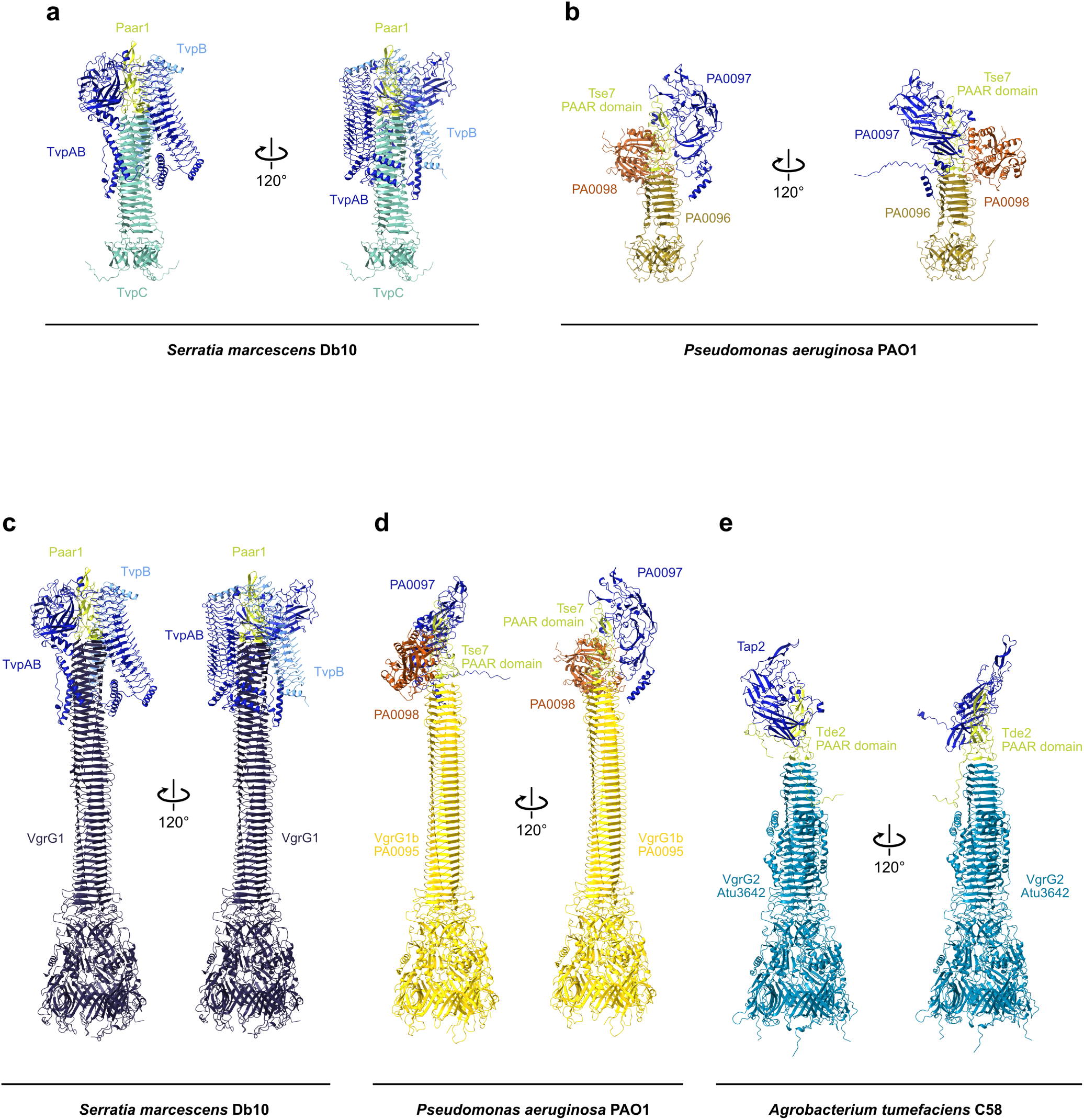
Predicted structures of accessory pre-complexes and VgrG-associated complexes of *Serratia marcescens*, *Pseudomonas aeruginosa* and *Agrobacterium tumefaciens*. **a)** Ribbon representation of the predicted structure of the *S. marcescens* Tvp accessory pre-complex. TvpAB is colored in medium blue, TvpB in light blue, TvpC in turquoise and Paar1 in light green. **b)** Ribbon representation of the predicted structure of the *P. aeruginosa* accessory pre-complex. PA0096 is colored in ginger, PA0097 in medium blue, PA0098 in rust and the PAAR domain of Tse7 effector in light green. **c)** Ribbon representation of the predicted structure of the *S. marcescens* full VgrG1-associated accessory complex. VgrG1 is colored in dark blue and accessory proteins are colored as in panel **a**. **d)** Ribbon representation of the predicted structure of *P. aeruginosa* full VgrG1b-associated accessory complex. VgrG1b is colored in gold and accessory proteins are colored as in panel **b**. **e)** Ribbon representation of the predicted structure of *A. tumefaciens* full VgrG2-associated accessory complex. VgrG2 is colored in sea blue, Tap2 in medium blue and the PAAR domain of Tde2 effector in light green. **a,b,e)** Additional representation and colouring according to the pLDDT score are provided in Supplementary Figures 8 and 11.

We then attempted to model the full VgrG1-associated complex from *S. marcescens* using AlphaFold2, but it was unable to generate a prediction. In contrast, AlphaFold3 successfully predicted a shorter version of this complex using only the tip of VgrG1 (aa 542-786), with a pLDDT score >75. We used this prediction as a basis to manually dock the accessory proteins onto our predicted structure of the full VgrG1 (**Figure 4**), allowing us to visualise how the full VgrG1 accessory complex could assemble (**Figure 6c**). The RMSD of the Cα atoms between the TvpC pre-complex (TvpC-TvpAB-TvpB-Paar1) and the TvpAB-TvpB-Paar1 complex with the tip of VgrG1 is only 0.907 Å between 798 pruned atom pairs, suggesting that only minor rearrangements occur during the transition between the pre-complex and the full-complex. Moreover, the binding of the accessory proteins to the tip of VgrG1 does not significantly modify the structure of the tip of VgrG1 (RMSD of 0.755 Å between 128 pruned atom pairs). We can thus expect that the tip of VgrG1 will simply be further stabilized/rigidified by the Tvp proteins compared to the protein alone (**Figure 4**) or in complex with only Paar1 (**Supplementary Figure 5b**).

Using the same approach of combining an AlphaFold3 intermediate prediction with manual docking, we were also able to reconstitute the full VgrG1b-associated accessory complex of *P. aeruginosa* (**Figure 6d**). In this case, AlphaFold3 produced a high confidence model of VgrG1b in complex with the PAAR domain of Tse7 (pLDDT >85) that served as a basis to dock PA0097 (TvpA) and PA0098 using the PAAR domain of Tse7 for the fitting. The presence of the PAAR domain of Tse7 on the VgrG1b tip does not change the overall structure of VgrG1b (RMSD of 0.683 Å between 560 pruned atom pairs). Furthermore, in this full VgrG1b-associated accessory complex, as in the pre-complex, neither PA0097 nor PA0098 directly interact with the β-prism of VgrG1b. Finally, we also obtained a structural prediction of VgrG2 from *A. tumefaciens* in complex with Tap2 and the PAAR domain of Tde2 using AlphaFold2 (pLDDT >85) (**Figure 6e** and **Supplementary Figure 11a-b**). In this complex, the DUF2169 of Tap2 wraps the tip of the PAAR domain of Tde2. Again, this interaction relies on hydrophobic regions of both Tap2 and the PAAR domain of Tde2, whilst Tap2 does not make any contact with VgrG2 (**Supplementary Figure 9c**). Overall, these structural predictions suggest that TvpA proteins containing only a DUF2169 domain do not directly interact with the VgrG tip. However, they can either act alone (*A. tumefaciens*) or in concert with an additional protein (*P. aeruginosa*) to chaperone the PAAR domain of effectors during the assembly of the T6SS machinery. In line with our Tvp nomenclature, we proposed to name the additional chaperoning protein in *P. aeruginosa* (PA0098, PRK06147) as TvpD.

### Classification of Tvp accessory systems

The diversity of Tvp systems means that phylogenetic classification is not straightforward. There are a number of components that are found in some systems but not in others, for example, TvpD or TvpB. Additionally, although a core conserved unit of VgrG-TvpA-Paar1 is a commonality, these individual proteins often contain divergent additional features and domains. VgrG sequences were found to be too variable to generate a useful phylogenetic classification, likely due to independent adaptations to facilitate species-specific interactions with baseplate components or effectors. For this reason, we compared the sequences and predicted structures of a number of PAAR and TvpA domains. A conserved sequence region corresponding to a shared structural arrangement was chosen for each protein (**Supplementary Figure 12**), and a phylogenetic tree was constructed using these TvpA-PAAR sequences concatenated (**Figure 7**). The analysis included proteins originating from bacterial species with an experimentally validated T6SS, including the *V. parahaemolyticus* system in which the DUF2169 (TvpA) protein was very recently characterised^10^. This tree further emphasizes the division of the different Tvp systems into three groups, with the *A. tumefaciens*, *P. aeruginosa* and *S. marcescens* systems serving as model representatives of each class (**Figure 7a**), consistent with the three structural assemblies described above. Also consistently, the *V. parahaemolyticus* system, whose DUF2169 gene (VP1398) is flanked by genes including one encoding a protein with a DUF6484 domain (VP1395), falls into the same group as *P. aeruginosa*, which also possesses a DUF6484-type TvpC protein.

**Figure 7.**
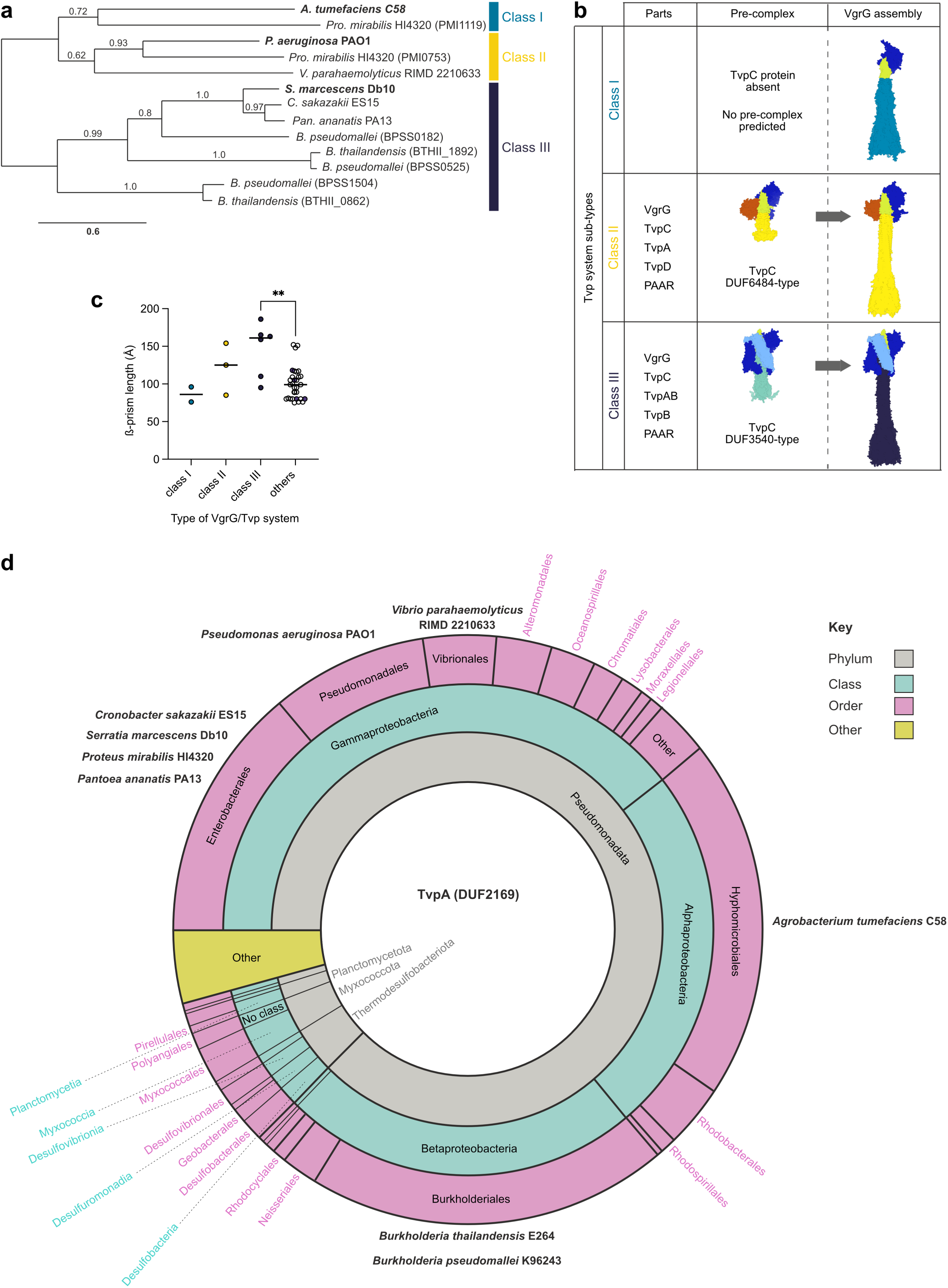
Classification of the Tvp system into Classes I, II and III based on structural, functional and phylogenetic comparisons. **a)** Phylogenetic tree based on PAAR and TvpA as core components of the Tvp systems, from selected bacteria with an experimentally validated T6SS. Where strains contain multiple Tvp systems, the accession number for the TvpA protein is enclosed in brackets to distinguish between them. The tree was constructed from conserved PAAR-TvpA sequences which were concatenated and aligned using MUSCLE^39,47^ (Supplementary Figure 12). **b)** Graphic table summarising the three different classes of Tvp systems. The table illustrates the structure and function relationships between the Tvp systems of Class I (light blue), Class II (yellow) and Class III (dark blue), represented by *A. tumefaciens* C58, *P. aeruginosa* PAO1 and *S. marcescens* Db10 respectively. **c)** Taxonomic distribution of TvpA (DUF2169) genes in available sequenced bacterial genomes. 4056 proteins with the InterPro entry IPR018683 (Pfam PF09937)^47,48^ are represented. Segments are weighted by number of species.

By combining our structural predictions of three representative assemblies and our phylogenetic analysis of *paar*-*tvpA*, we can define a functional classification for the Tvp accessory complexes (**Figure 7b**): The simplest system, consisting of VgrG, PAAR and TvpA, is described as Class I and is represented by *A. tumefaciens* C58^7^. Class II, represented by *P. aeruginosa* PAO1, is distinguished by a number of additional proteins which we predict, based on our observations in *S. marcescens*, will form a pre-complex using a short TvpC trimer protein (DUF6484) which is a partial structural mimic of VgrG with an OB-fold and very short β-prism. Class II systems have the unique property of having an additional predicted PAAR chaperone which we call TvpD. Class III systems, represented by our *S. marcescens* Db10 system, have a longer TvpC protein (DUF3540) and contain an additional unique component, the pentapeptide-repeat domain (PPR) containing protein TvpB, which is included as a C-terminal extension of TvpA and an independent protein. This classification suggests an evolutionary trend towards increased complexity in the set of accessory proteins and in the steps involved in T6SS assembly.

As noted above, VgrG1 in *S. marcescens* possesses an exceptionally long β-prism (**Figure 4a**). Therefore, we performed a comparison with VgrG proteins from Tvp-containing, experimentally validated T6SS clusters, as well as other VgrG proteins encoded elsewhere within the corresponding genomes, to assess whether the length of the β-prism correlates with the number of accessory proteins. To this end, all VgrG structures were predicted with AlphaFold3 and the length of the β-prism was measured manually whenever possible (**Supplementary Table 5**). Among the 45 VgrG predictions obtained, we excluded VgrG4a and VgrG4b from both *Burkholderia* strains from our analysis due to inaccurate folding predictions of the β-prism tip. Despite the small sample size of Class I and Class II VgrG/Tvp systems, this measurement shows that, overall, VgrG proteins belonging to the Tvp Class III systems have a longer β-prism than other VgrG proteins and this difference is significant (p-value < 0.05), at least compared to VgrG proteins which do not function with Tvp accessory proteins (**Figure 7c**). However, given the distribution of β-prism lengths within each class, this is likely not the sole parameter driving the diversification of Tvp proteins.

Since TvpA is the common and defining features of the three Tvp classes, we analysed the taxonomic distribution of Tvp systems by screening available sequenced bacterial genomes for the presence of *tvpA* (DUF2169) (**Figure 7d**). This analysis showed that TvpA, which is classified as IPR018683 on the InterPro database and represented by 4056 protein sequences, is widespread and found in a number of clinically and environmentally important bacterial species. Most sequences belong to the phylum *Pseudomonodata* but there are a number of other phyla represented (*Acidobacteriota, Myxococcota, Planctomycetota, Thermodesulfobacteriota*). Moreover, to validate our classification, we carried out an additional phylogenetic analysis using a more diverse set of concatenated sequences from the other bacterial phyla (**Supplementary Figure 13**). This showed that these diverse sequences accommodate perfectly into the same three functional and structural Tvp classes we defined, strengthening our classification (**Supplementary Figure 14a-b**).

## DISCUSSION

In this study, we have defined a broad family of T6SS accessory complexes which all involve DUF2169-containing proteins and are subdivided into three classes, two of which mediate a previously-undescribed two-step assembly process. Detailed characterisation of the Class III Tvp system of *S. marcescens* revealed that a TvpC-containing pre-complex templates the formation of the subsequent VgrG1-containing complex, which supports overall T6SS function and allows delivery of an unusual VgrG1-associated membrane-depolarising effector.

The T6SS is widely used by Gram-negative bacteria to deliver toxic antibacterial effector proteins into neighbouring rivals. Such effectors are encoded next to a partner gene encoding the cognate immunity protein, which provides the producing cell with protection against the effector, whether self-produced or delivered by a neighbouring cell. Here we characterized Vai1, a new immunity protein in *S. marcescens*, which provides protection against an effector delivered through the VgrG1 pathway. Intriguingly, we were unable to formally identify its cognate effector using standard approaches, although an interaction with VgrG1 was observed by native co-purification. Since this interaction was observed only when using Vai1 as bait under conditions where T6SS-mediated protein delivery between cells can take place, it is likely to occur exclusively in target cells and not during the assembly of the machinery where the VgrG1 tip will be shielded by accessory proteins. Vai1 is encoded immediately downstream of the *vgrG1* gene, the first gene of the T6SS cluster, suggesting that VgrG1 itself is likely to be the cognate effector. This idea is supported by the fact that the gene downstream of *vai1* does not encode an effector, the absence of effector candidates alongside Vai1 homologues in other genomes, and the lack of involvement of the other known T6SS effectors. However, despite having a longer C-terminus than VgrG2, no classical toxin domain could be identified in VgrG1 and this extension appears to form an elongation of the structural β-prism, making it unclear how VgrG1 exerts toxicity. An attractive hypothesis is that the C-terminus contains a cryptic toxic peptide, which would be toxic only when cleaved off from the full structural protein. Indeed, the last 26 residues at the C-terminus are largely hydrophobic and an antimicrobial peptide prediction calculator^17^ indicates that this region could exist as an α-helix with ten hydrophobic amino acids on the same face, when not part of the full VgrG1 structure. In this model, short amphipathic α-helices, released from intact VgrG1 proteins, would interact with the lipids of the target cell membrane and cause the membrane depolarization we observed in intoxicated cells. The mechanism by which this putative antimicrobial peptide would be released from the VgrG1 protein is yet to be determined, although the Tvp accessory complex is likely to prevent it occurring prior to secretion and Vai1 may prevent it occuring in resistant cells. Another, more unlikely, possibility is that the VgrG1-Paar1 spike forms a “mechanical toxin” similar to R-type tailocins from *P. aeruginosa,* whose physical action on the inner membrane induces depolarization^18^. Resistance to this type of toxin is usually mediated through modification of the lipopolysaccharide receptor, rather than through the presence of a cognate immunity protein^19,20^. Thus, if VgrG1-Paar1 forms such a mechanical toxin, it remains unclear how Vai1 could intercept the incoming spike and prevent its toxic activity.

Loading of effectors into the T6SS machinery often involves accessory proteins, which can act as chaperones or adaptors to stabilise or guide them to the core machinery. DUF2169 proteins represent examples of such accessory proteins, and have been shown to be required for the delivery of Tde2 in *A. tumefaciens*^7^, and a PAAR-like DUF4150-containing effector in *V. parahaemolyticus*^10^. Here, we show that TvpAB, a DUF2169-containing protein, acts in concert with TvpB and TvpC to load the Paar1 protein of *S. marcescens* into the T6SS machinery by forming a pre-complex, where TvpC is subsequently replaced by VgrG1 in the full T6SS assembly. Similar and also simpler versions of these Tvp accessory proteins exist in a large number of bacteria and all of these accessory complexes share the presence of a DUF2169-containing protein that we call TvpA. Indeed, based on extensive structural predictions reinforced by phylogenetic analysis, we propose a classification of Tvp systems into three distinct classes, built upon a progressively increasing complexity in terms of both accessory components and assembly steps. Class I, exemplified by *A*. *tumefaciens*, is the simplest version, with a single accessory protein, TvpA, which stabilizes the PAAR domain of Tde2 directly onto VgrG2. Class II, represented by *P. aeruginosa*, also includes a TvpA protein, which we predict will form a pre-complex with a TvpC-like protein and interact with the PAAR domain of the Tse7 effector before loading it onto the tip of VgrG1b. This class also contains another protein that we call TvpD, which could be classified as a co-chaperone for the PAAR domain. This aligns with a bioinformatic paper which speculates that PRK06147 may represent a novel T6SS adaptor^21^. In both Class I and Class II systems, the accessory proteins are in close interaction with the PAAR domains of the effectors while barely interacting with VgrG proteins. Finally, Class III is the most sophisticated assembly and is represented by the *S. marcescens* Tvp system. Overall, this classification aligns with the three synteny categories described by Sachar *et al.*^10^, although we provide additional detail on the different groups, especially in terms of assembly. Our structural prediction for the TvpA domain of *S. marcescens* Db10 is very similar to the crystal structure of the *V. xiamenensis* DUF2169 protein, aligning with an RMSD of 1.076 Å (**Supplementary Figure 15**), confirming the accuracy of the prediction. Both *V. xiamenensis* and *V. parahaemolyticus* encode a DUF6484 protein, a hallmark of the Class II Tvp system, thus we can predict that T6SS assembly in these organisms will require the formation of a pre-complex similar to the one we modelled for *P. aeruginosa* (**Figure 6b**). In addition, VP1399 in *V. parahaemolyticus*, which was not functionally characterized by Sachar *et al.*^10^, likely encodes a TvpD protein chaperoning the PAAR domain of the VP1415 toxin, in association with the TvpA protein, VP1398.

Our analysis indicates that the DUF2169 domains of TvpA (*A. tumefaciens/P. aeruginosa*) and TvpAB proteins (*S. marcescens*) strongly interact with the PAAR domains in all three classes, making this interaction a common structural feature of the different Tvp systems. This interaction seems to serve mainly to protect hydrophobic regions in PAAR domains, confirming that DUF2169 domain is a PAAR chaperone. However, the extent of the hydrophobic regions in PAAR proteins differs between the three systems. This results in a different orientation of the PAAR-TvpA tandem within the pre-complexes (when existing) or within the full complexes with VgrG proteins. Indeed, when structural models are superimposed by their PAAR domain, a moderate rotation of the DUF2169 is observed between Class I and Class II (∼10°) while this rotation reaches almost 45° for Class III (**Supplementary Figure 16a**). When models are superimposed based on the short α-helix in a protruding loop of the DUF2169 domain which is important for the interaction with the hydrophobic patch of PAAR^10^, again the β-sheets of the DUF2169 domain move by ∼10° in Class II and by ∼25° in Class III compared to Class I (**Supplementary Figure 16b**). This highlights the ability of the DUF2169 domain to adapt its chaperoning effect to the biophysical properties of its cognate PAAR domain.

Our findings that TvpC and VgrG1 form two mutually exclusive complexes (**Figure 3**) provide new insights into the assembly in the T6SS machinery and open new questions. Why do some T6SS require the formation of pre-complexes (Tvp systems Classes II and III) while others do not (Tvp system Class I, and VgrG with non-Tvp accessory proteins)? We could hypothesize that, in addition to their stabilizing effect on PAAR protein, Tvp proteins might interact with the baseplate and participate in its docking to the membrane complex, given that VgrG1 serves as the central hub of this baseplate. Indeed, it has been shown that both VgrG and PAAR are required for the proper localization of the baseplate during T6SS assembly^22,23^. Two key characteristics of *S. marcescens* Paar1 and VgrG1 components may implicate the pre-complex in the recruitment of baseplate components. First, Paar1 is a small protein, with a high degree of hydrophobicity and devoid of any other domains. Stability and interaction possibilities for Paar1 during T6SS assembly would thus be limited without the help of accessory proteins. Secondly, VgrG1 has a remarkably long β-prism. Such length means that VgrG1-Paar1 tip (estimated at 242 Å) will penetrate deeper into the cavity of the membrane complex and might disturb its closed resting state which is important for the proper assembly of the T6SS machinery^24^. Thus, having a pre-complex with a VgrG-like protein, namely TvpC, harbouring a smaller, more typical β-prism would ensure the correct recruitment of the baseplate without compromising the integrity of the membrane complex. During initial studies, we looked for any interaction between Tvp accessory proteins and baseplate components using bacterial two-hybrid assays. We observed an interaction only between TssK and TvpC (**Supplementary Figure 17a,b,d**). Moreover, TssK also interacts with VgrG2 but not with VgrG1 (**Supplementary Figure 17c-d**). These data suggest that both TvpC and VgrG2 could participate in the recruitment of baseplate components during the assembly of the T6SS apparatus. Cryo-EM structures of baseplates from different organisms instead highlighted an interaction between VgrG and TssF wedges^22,25,26^. However, the absence of a Gp27 domain in TvpC and the high flexibility of TssK^26^ could explain variations in the overall shape and assembly of the T6SS baseplate between systems^25^. The Gp27 domain of VgrG is required to promote Hcp tube polymerisation^27^. Therefore, as TvpC does not possess a Gp27 domain, the T6SS machinery could not fully assemble on the pre-complex, meaning that TvpC has to be replaced by VgrG1 to allow Hcp polymerisation, concomitant sheath assembly and, ultimately, firing. The mechanism of this sequential assembly and replacement of TvpC with VgrG1 remains to be fully elucidated in future studies.

The cryo-EM reconstruction of the *V. cholerae* baseplate highlighted the presence of a cavity formed by the baseplate periphery (TssK oligomers) around the central spike. Such a cavity was proposed to accommodate effectors associated with VgrG or PAAR^25^. Moreover, Liang *et al*., demonstrated that some T6SSs have an “onboard checking” mechanism by which the presence of effectors loaded on the machinery is a prerequisite for T6SS secretion^28^. In the absence of specific effector proteins or domains delivered by VgrG1^14^, this cavity could instead be filled with the Tvp accessory proteins in order to allow stable baseplate assembly and/or fulfil such an assembly checkpoint prior to firing. This hypothesis could explain why the VgrG1 pathway, without any effectors decorating the spike, is still able to assemble, fire and deliver Hcp-dependent effectors^14^. In contrast, VgrG2 utilises large specialised PAAR proteins, Rhs1 and Rhs2. More generally, from our set of representative strains, it seems that Class III Tvp systems are found with small PAAR proteins without effector domains, whereas Class I and II systems are found with PAAR proteins containing specialised effector domains. Thus, the requirement for TvpAB-TvpB proteins could be linked to small PAAR proteins and a resulting need to occupy the baseplate cavity. In the *P. aeruginosa* Tvp system, which also forms a pre-complex, the PAAR domain is part of the Tse7 effector, thus the toxic moiety could partially fill the cavity, explaining why smaller Tvp proteins (TvpA and TvpD) would be required.

Collectively, our findings identify a new set of T6SS accessory proteins, named Tvp, which are associated with DUF4150-type PAAR proteins across many organisms and are required for the assembly and delivery of the corresponding VgrG-PAAR-containing puncturing structure with its payload of effectors. We have defined three classes of Tvp system, which display increasing complexity in their composition and assembly. This variation likely reflects specific requirements for T6SS assembly, such as a need to accommodate the length of the VgrG β-prism and occupy space in the baseplate cavity not filled by effector domains. Our findings also suggest the existence of a new type of VgrG effector which is able to induce loss of membrane potential in target cells in the absence of a separate effector domain. It is clear that the basic mechanism of the T6SS is highly versatile, with accessory proteins and complexes able to fine-tune and adapt its function.

## METHODS

### Bacterial strains, plasmids and culture conditions

The strains and plasmids used in this study are listed in **Supplementary Tables 1-2**. Genetic variants of *S. marcescens* Db10 were generated by allelic exchange using the suicide plasmid pKNG101 as previously described^12^. Streptomycin-resistant derivatives of Db10 strains were generated by introducing a SNP in *rpsL* which resulted in asparagine (N) being encoded at amino acid position 43. This SNP was either introduced by phage ΦIF3-mediated transduction of the resistance allele from *S. marcescens* Db11 or by recombineering using a modified version of the pORTMAGE plasmid (pSC3048) in conjunction with oligonucleotide CE75 carrying the desired SNP^29^. Plasmid-based constitutive expression of genes for complementation of gene deletions was achieved using the pSUPROM plasmid.

Unless otherwise stated, bacterial cultures were grown in LB (10 g.L^-1^ tryptone, 5 g.L^-1^ yeast extract, 10 g.L^-1^ NaCl, with 1.8 g.L^-1^ agar for solid media) at 37 °C for *E. coli* and 30 °C for *S. marcescens* and *P. fluorescens*. When required, media were supplemented with antibiotics: carbenicillin (Ap) 100 μg.mL^-1^, chloramphenicol (Cm) 25 μg.mL^-1^ kanamycin (Kn) 100 μg.mL^-1^ and streptomycin (Sm) 100 μg.mL^-1^. For measurement of Hcp1 secretion by immunoblot, *S. marcescens* Db10 was grown in low salt liquid LB at 30 °C (10 g.L^-1^ tryptone, 5 g.L^-1^ yeast extract, 5 g.L^-1^ NaCl).

### Anti-bacterial co-culture assay to assess T6SS-mediated activity

We utilised our standard anti-bacterial co-culture assay^12^. Attacker and Sm-resistant target strains of *S. marcescens* Db10 and its derivatives, *S. marcescens* SM39, or *P. fluorescens* 55 (Sm-resistant), were patched from a single colony and grown overnight on LB agar. Bacterial cells were recovered from the patches with a sterile loop and suspended in 1 mL of LB broth. Cell suspensions were normalised to an OD_600nm_ of 0.5 and attackers and targets mixed at a 1:1 ratio. 25 μL of the mixture was spotted onto solid LB and grown for 4 h (for inter-species assays) or 7.5 h (for intra-species assays). Following the co-culture, cells were recovered in 1 mL of LB broth and the number of surviving target cells was enumerated by serial dilution, plating onto LB agar supplemented with Sm and determining viable counts.

### Membrane potential and membrane permeability analysis

A 25 μL mixture of attacker and target strains of *S. marcescens* Db10 were co-cultured at an initial ratio of 1:2 (attacker:target) on solid LB media for 4 h at 30 °C, then cells were recovered and suspended in 1× phosphate-buffered saline (PBS) at 10^6^ cells/mL. Both DiBAC_4_(3) (Bis-[1,3-Dibutylbarbituric Acid] Trimethine Oxonol; Thermo) at 10 μM final concentration and propidium iodide at 1.5 μM final concentration were added simultaneously to each cell suspension, followed by incubation in the dark for 30 min. As a control, a cell suspension, derived from growth of a single colony on solid media, of the target strain DWL08 Δ*SMDB11_2245-46* Δ*tssE* Sm^res^ was used to inoculate a 5 mL culture of LB low salt broth at a starting OD_600nm_ of 0.05 and cells were grown for 2.5 h at 30 °C, to reach an OD_600nm_ of ∼0.5 (mid-log growth phase). Cells equivalent to 1 mL of cells at an OD_600nm_ of 0.1 were collected by centrifugation. The cell pellet was re-suspended in 1 mL of PBS supplemented with melittin at 0.02 mg.mL^-1^ (final concentration of 6.32 μM) and incubated with agitation at 30 °C for 2 h, prior to staining. After staining with DiBAC_4_(3) and propidium iodide, cells were directly analysed in a FACS LRS Fortessa equipped with 488 nm and 561 nm lasers (Becton Dickinson), using thresholds on side and forward scatter to exclude electronic noise. Channels used were Alexa 488 (Ex 488 nm, Em 530/30 nm) for DiBAC_4_(3) and Alexa 568 (Ex 561 nm, Em 610/20 nm) for propidium iodide. All bacterial suspensions were normalised to ∼10^6^ cells/mL prior to analysis. Analysis was performed using FlowJo v10.4.2 (Becton Dickinson).

### Immunodetection of cellular and secreted proteins

*S. marcescens* Db10 and derivates were grown, aerated, for 5 h at 30 °C in 50 mL of low salt LB broth. Hcp1 detection was carried out on cellular and secreted fractions. Cellular protein samples were obtained by collecting 100 μL of culture from which cells were recovered by centrifugation. The cell pellet was resuspended in 100 μL of 2 x SDS sample buffer (100 mM Tris-HCl pH 6.8, 3.2 % SDS, 3.2 mM EDTA, 16% glycerol, 0.2 mg.mL^-1^ bromophenol blue, 2.5 % β-mercaptoethanol). Secreted protein samples were generated by combining 100 μL of culture supernatant with 100 μL 2 x SDS sample buffer. An amount equivalent to 1 mL of cells at an OD_600nm_ of 0.015 was loaded for the cellular fraction and an amount equivalent of supernatant generated from 1 mL of culture at an OD_600nm_ of 0.015 was loaded for the secreted fraction. Hcp1 was detected using anti-Hcp1 rabbit primary antibody at a dilution of 1:6000 and horseradish peroxidase (HRP)-conjugated anti-rabbit secondary antibody (Bio-Rad no:170-6515) used at a dilution of 1:10,000.

### Immunoprecipitation of epitope-tagged proteins with magnetic beads

Derivatives of *S. marcescens* Db10 encoding epitope tagged versions of each protein of interest at the normal chromosomal location were generated by replacement of the wild type allele of the gene. Bacterial cultures were inoculated at a starting OD_600nm_ of 0.025 and grown for 5 h in 50 mL of LB low salt broth to a final OD_600nm_ of ∼2-2.5. Cells were recovered by centrifugation at 48,400 x g for 20 min, 4 °C. The cell pellet, equivalent to 50 mL of culture, was re-suspended in 1 mL of ice-cold lysis buffer (Tris pH7.5 20 mM, NaCl 150 mM, EDTA 0.5 mM, Triton X-100 0.1% + Roche complete EDTA-free protease inhibitor cocktail). Lysis was carried out on ice using sonication with a 1.6 mm microtip probe delivering 30 % amplitude for 8 x 15 sec with 30 seconds off between each pulse. Cell debris was removed by centrifugation at 20,000 g and the clarified lysate was incubated with 30 μL of magnetic anti-HA (Rabbit mAb bead conjugate from New England Biolabs, Ref. 11846S) or anti-FLAG (M2 magnetic beads from Merck, Ref. M8823) beads for 1-2 h at 4 °C, 40 rpm. Beads were washed with 1 mL of lysis buffer 3 times, with a further wash carried out with lysis buffer which lacked Triton X-100. For elution the beads were incubated with an SDS-based buffer (5% SDS, 50 mM Tris pH 7.5, 150 mM NaCl) for 5 min at 70 °C; the beads were then removed using a magnetic separation rack.

### Small scale affinity purification of His-tagged proteins with magnetic NTA-beads

Strain construction, growth conditions and protein isolation procedures were the same for small scale affinity purification (pull-down) as for the immunoprecipitation of epitope-tagged proteins, with the exception of the lysis and wash buffer chemistry. Lysis was carried out using a Tris-HCl 50 mM, NaCl 100 mM, Imidazole 20 mM + Triton X-100 0.5 % lysis buffer with the sonication settings described above. The clarified lysate was incubated with 30 μL of Ni-NTA magnetic agarose beads (Qiagen Ref. 36113) for 1-2 h at 4 °C, 40 rpm. The wash buffer used was the same as the lysis buffer but with the inclusion of 50 mM imidazole. Elution was using the same SDS-based elution buffer as above.

### Immunoprecipitation from cells grown on solid media

To determine interaction partners of the immunity protein Vai1 (SMDB11_2245), strains were grown on solid media, similar to the anti-bacterial co-culture assay, to permit T6SS delivery of effectors between cells. In this case the strain was self-delivering the VgrG1-dependent effector. A single colony of each strain was patched and grown overnight on LB agar. Bacterial cells were recovered from the patches with a sterile loop and suspended in 1 mL of LB broth. Suspensions were normalised to 500 μL of OD_600nm_= 2. The suspension was spotted as 10 x 50 µL spots on a single LB agar plate and grown for 5 h at 30°C and the spots were harvested in 10 mL of LB from the plate which gave an OD_600nm_ of ∼2. From this point onwards, the protocol was exactly the same as for immunoprecipitation with anti-FLAG M2 magnetic beads above.

### Mass spectrometry and label-free quantitation

Eluted protein samples in SDS elution buffer (5% SDS, 50 mM Tris pH 7.5, 150 mM NaCl) were analysed by the ‘FingerPrints’ Proteomics Facility, University of Dundee. In brief, protein samples were subjected to on-column tryptic digestion and clean-up using S-Trap columns before being analysed by nLC-MS/MS using a Q-Exactive^TM^ Plus mass spectrometer (Thermo Fisher Scientific) with 120 min nLC gradient. All samples were analysed in biological triplicate. Label-free quantitation was performed using MaxQuant^30^ and comparative data analysis performed using Perseus^31^. Proteins were considered to be significantly enriched in abundance in test (epitope-tagged) samples compared with control samples if LFQ intensity was significantly increased according to the criteria: log2 fold change test/control > 2, p < 0.05. Proteins were considered to be present in test and absent in control samples if detected in all three replicates of the test (with log2 LFQ intensity >25) and either not detected in any replicate of the control or in only one replicate of the control (when that single replicate gives log2FC test/control > 3).

### Bacterial two-hybrid assay

Bacterial two-hybrid analyses were performed following established protocols^32,33^. *E. coli* MG1655 Δ*cyaA* was co-transformed with combinations of a pUT18-based and a pT25-based plasmid and the color of the resulting transformants scored on MacConkey media with Ap, Cm and 0.2% maltose (positive result being red). For quantitative measurement of the interaction, β-galactosidase assays were performed as described^12^ on double-transformed MG1655 Δ*cyaA* grown at 30°C in LB and permeabilized with toluene. Replicate assays were performed on three independent transformants.

### Structural prediction and analysis

Protein and protein complex structural predictions were generated using AlphaFold2 and AlphaFold2 multimer v. 3 (hosted on the University of Dundee HPC cluster and implemented with ColabFold), or using AlphaFold3 (hosted on https://alphafoldserver.com) ^34–36^. Figures depicting AlphaFold predictions were prepared with ChimeraX^37^. Structural predictions were superimposed using the Matchmaker tool of ChimeraX, with the chain-pairing method when required, and with default fitting parameters. Structural comparisons were made using the structural alignment function in PyMol (The PyMOL Molecular Graphics System, Version 3.0 Schrödinger, LLC). The transmembrane domain of Vai1 was identified using TMHMM 2.0^38^.

### Phylogeny and taxonomy of Tvp systems

Multiple sequence alignments of concatenated TvpA and DUF4150 (PAAR) amino acid sequences were produced using the MUSCLE algorithm^39^, and coloured with ClustalX^40^. Alignment figures were prepared in the Jalview platform^41,42^. The MUSCLE alignment was used for the generation of a phylogenetic tree. This was computed using phylogeny.fr^43^ with PhyML^44^ using a WAG substitution model^45^ and 500 bootstrap replicates. The output tree was visualised using TreeDyn^46^. The taxonomic distribution of the TvpA protein was adapted from the InterPro^48^ taxonomy graphic available for the DUF2169 classification entry (IPR018683).

### Data Availability

All data supporting the findings of this study are available within the paper and its supplementary information files.

## ACKNOWLEDGMENTS

This work was supported by Wellcome [104556/Z/14/Z, Senior Research Fellowship, and 220321/Z/20/Z, Senior Research Fellowship Renewal, held by SJC]. Mass spectrometry was performed by the FingerPrints Proteomics Facility at the University of Dundee, supported by a Wellcome Technology Platform Award [097945/B/11/Z]. We thank the Flow Cytometry and Cell Sorting Facility, the High Performance Computing Facility and Dr James Abbott at the University of Dundee for access to facilities and expert assistance. We thank Matthias Trost and members of his lab, particularly Julien Peltier and Akshada Gajbhiye, for mass spectrometry analysis of preliminary pulldown and immunoprecipitation experiments and advice on sample preparation. We would also like to acknowledge Duncan Lockie, Deborah Pasquali and Yi-Chia Liu for the construction of several mutant strains used in the study and Sebastien Santini (CNRS/AMU IGS UMR7256) and the PACA Bioinfo platform for the availability and management of the phylogeny.fr website. For the purpose of Open Access, the authors have applied a CC BY public copyright licence to any Author Accepted Manuscript version arising from this submission.

## AUTHOR CONTRIBUTIONS

C.E., L.M. and S.J.C. conceived the study, designed experiments and analysed data; C.E. and L.M. performed experimental work; C.E. and L.M performed bioinformatics and structural analyses; C.E., L.M. and S.J.C. prepared the figures, and L.M. wrote the manuscript with contributions from C.E. and S.J.C.

