## Supplementary Data for "Tvp complexes support formation of the VgrG-PAAR spike during Type VI secretion system assembly"

#### Supplementary Figure 1

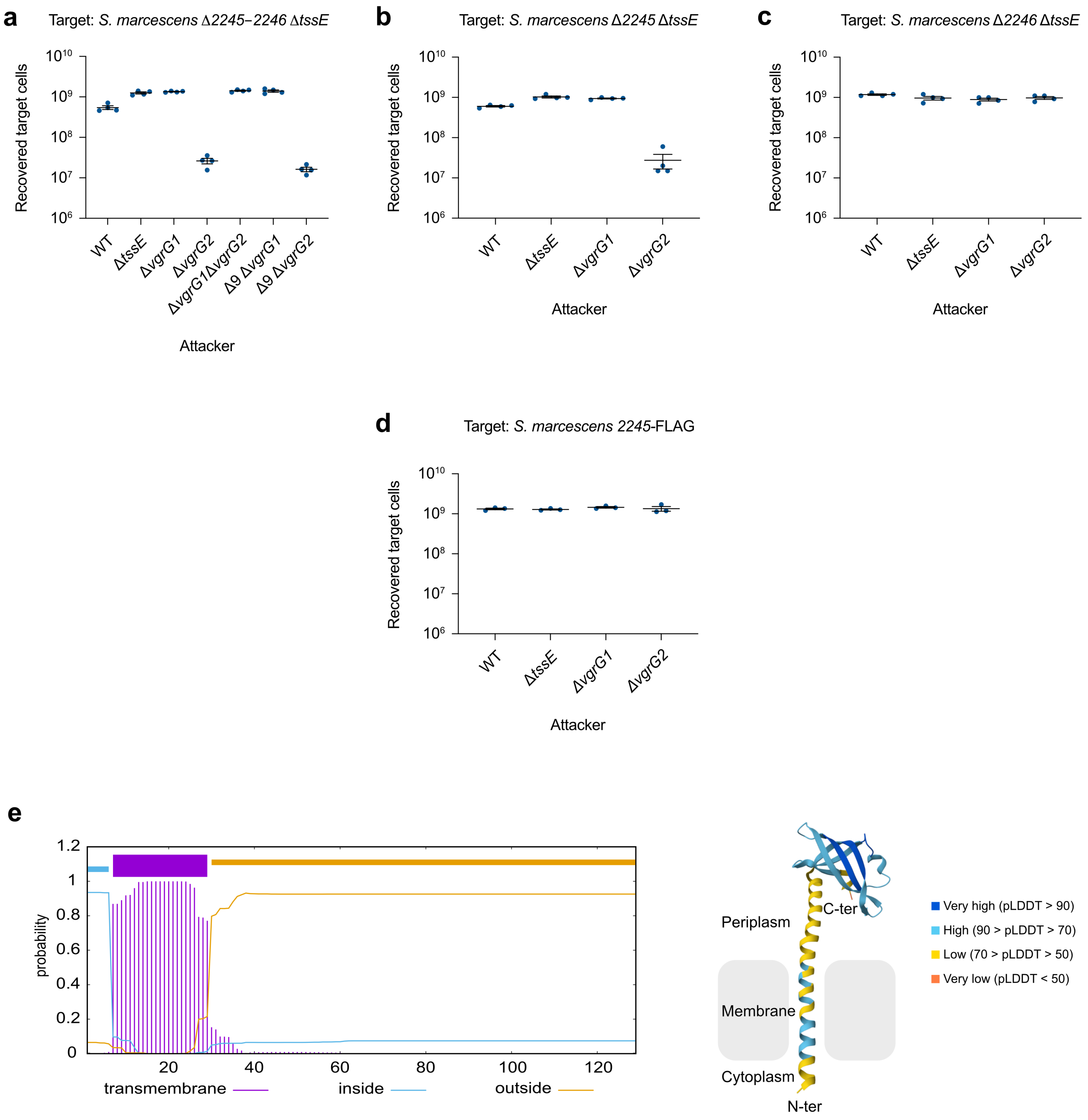


**Supplementary Figure 1. Vai1 is a new immunity protein conferring protection against the effector delivered by the VgrG1 pathway of *Serratia marcescens* Db10 a)** Recovery of *S. marcescens* Db10 Δ*2245-2246* Δ*tssE* target cells following co-culture with wild type (WT) and mutant strains of *S. marcescens* Db10. **b)** Recovery of Db10 Δ*2245* Δ*tssE* target cells following co-culture with WT and mutant strains of Db10. **c)** Recovery of Db10 Δ*2246* Δ*tssE* target cells following co-culture with WT and mutant strains of Db10. **d)** Recovery of the Db10 Δ*2245-2246* Δ*tssE* target complemented with *2245*-FLAG expressed from plasmid pSUPROM, following co-culture with WT and mutant strains of Db10. In **a**-**d**, co-cultures were performed with an initial ratio of 1:1. **e)** Left, Vai1 transmembrane helices prediction using TMHMM 2.0. The potential transmembrane helix in Vai1 (amino acids 7-29) is highlighted in purple. Right, Ribbon representation of the Vai1 structure predicted by AlphaFold2. The protein is coloured based on the reported confidence of the AlphaFold modelling, from orange (pLDDT <50, very low confidence) to dark blue (pLDDT >90, very high confidence). Vai1 insertion in the inner membrane is depicted based on the TMHMM prediction. **a-d)** Data sets are displayed as mean +/- SEM (n=4) with individual data points overlaid.

### Supplementary Figure 2

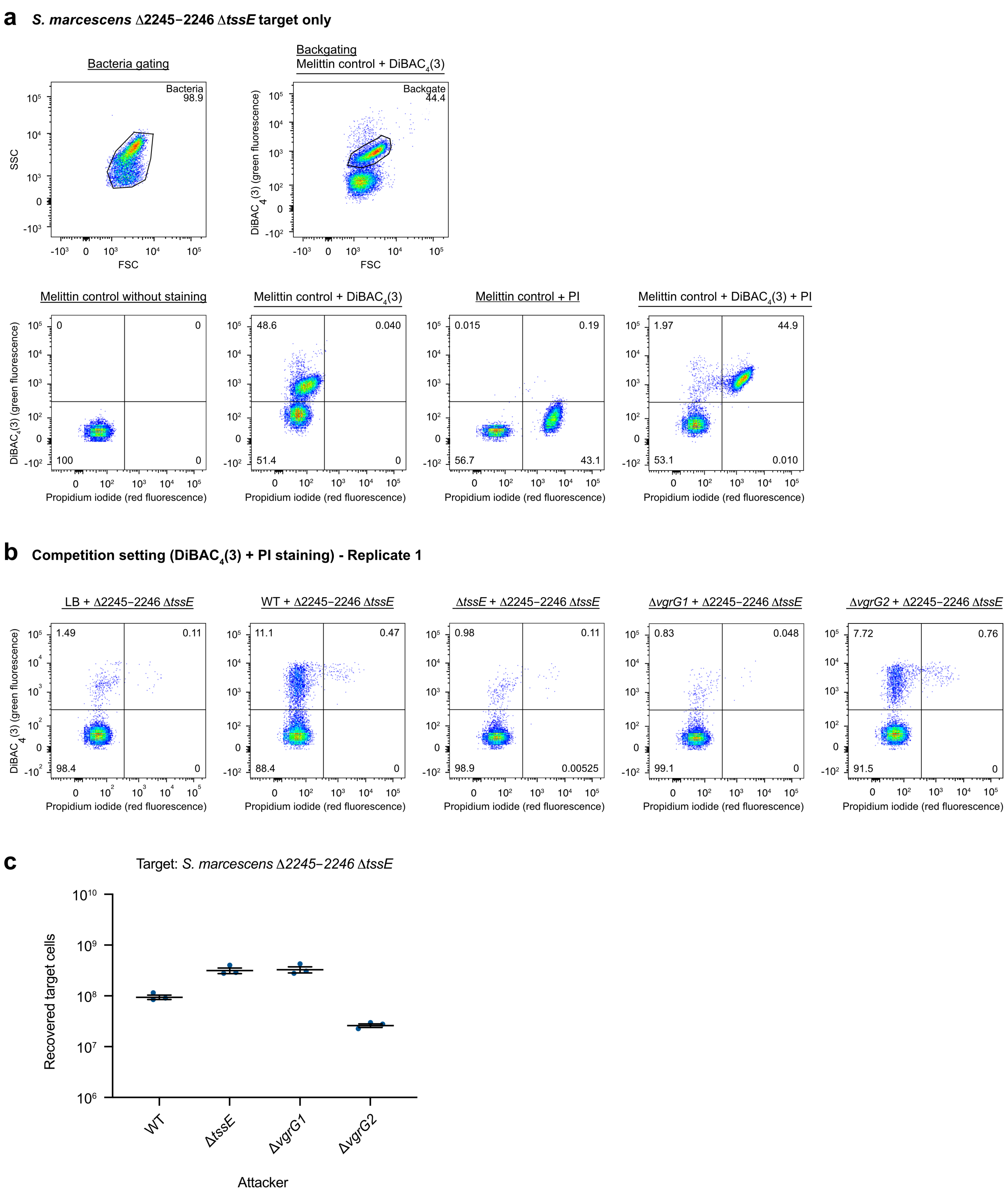


**Supplementary Figure 2. Example of flow cytometry experiments from the analysis presented in Figure 1g. a)** Analysis of *Serratia marcescens* Db10 *Δ2245-2246 ΔtssE* target only. Top panels illustrate the gating strategy. Left: Bacterial cells were selected using side scatter (SSC) vs. forward scatter (FSC). Right: Bacterial cells were treated with melittin, stained with $DiBAC_4(3)$ (green fluorescence) and were selected using $DiBAC_4(3)$ fluorescence vs. FSC. Bottom panels illustrate the quadrant strategy. Cells subjected to melittin treatment and stained with $DiBAC_4(3)$ only define the depolarised cells quadrant (green fluorescence) while those treated with propidium iodide only were used to define the permeabilized cells quadrant (red fluorescence). Cells treated with melittin and stained with both $DiBAC_4(3)$ and propidium iodide were used to establish the quadrant for cells that are simultaneously depolarised and permeabilized (green and red fluorescence). **b)** Analysis of bacteria following co-culture of *S. marcescens* Δ*2245-2246* Δ*tssE* target with wild type (WT) or mutant strains of Db10 for four hours at an initial attacker:target ratio of 1:2. After four hours of co-culture, cells were treated with both $DiBAC_4(3)$ and propidium iodide. Data shown correspond to one of the four replicates presented in Figure 1g. **c)** Recovery of the *S. marcescens* Δ*2245-2246* Δ*tssE* target following co-culture with WT or mutant strains under the conditions described in panel **b**. Data sets are displayed as mean +/- SEM (n=3) with individual data points overlaid.

#### Supplementary Figure 3

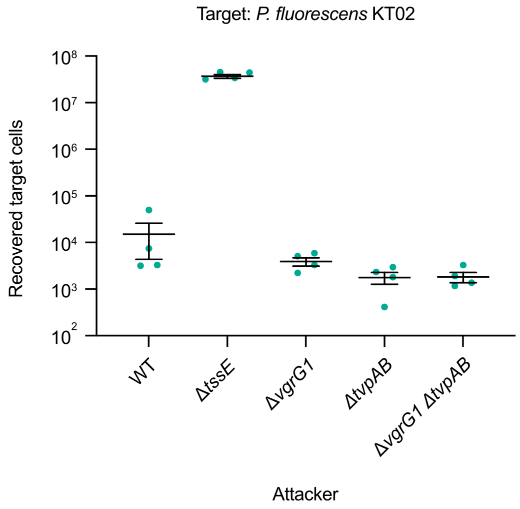


**Supplementary Figure 3. TvpAB exerts a repressive effect on the VgrG2 pathway, independently of VgrG1.** Recovery of *Pseudomonas fluorescens* target cells following co-culture with wild type (WT) and mutant strains of *Serratia marcescens* Db10 at an initial ratio of 1:1. Data sets are displayed as mean +/- SEM (n=4) with individual data points overlaid.

#### Supplementary Figure 4

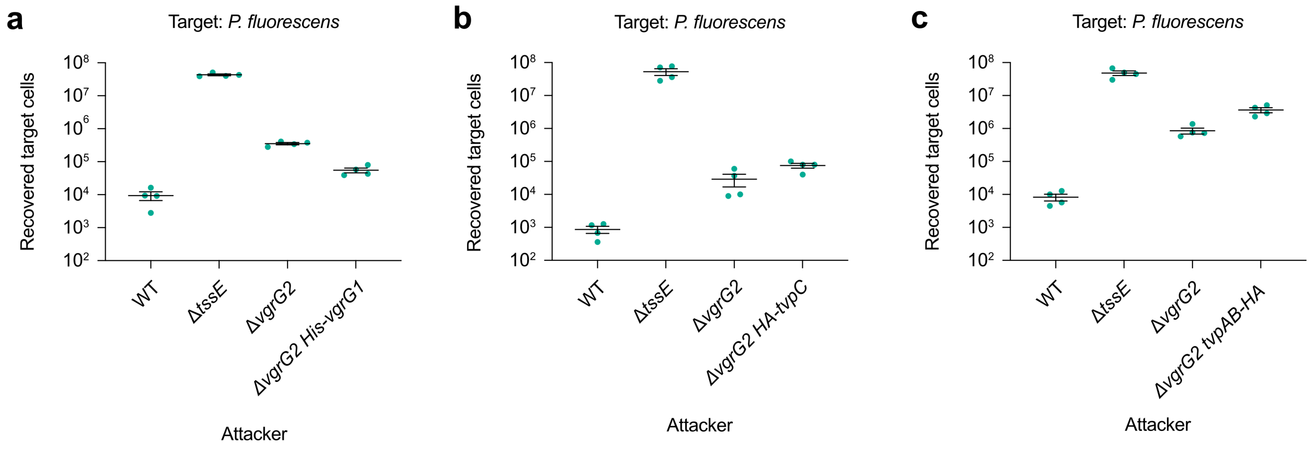


**Supplementary Figure 4. Epitope-tagged proteins used as bait in affinity purification and immunoprecipitation experiments retain function. a-c)** Recovery of *Pseudomonas fluorescens* target cells following co-culture with wild type (WT) and mutant strains of *Serratia marcescens* Db10 at an initial ratio of 1:1 for four hours. The functionality of the tagged proteins, namely His-VgrG1 **(a)**, HA-TvpC **(b)**, and TvpAB-HA **(c)**, was assessed in a Δ*vgrG2* background where loss of function in the VgrG1 pathway results in a complete loss of T6SS activity. Data sets are displayed as mean +/- SEM (n=4) with individual data points overlaid.

### Supplementary Figure 5

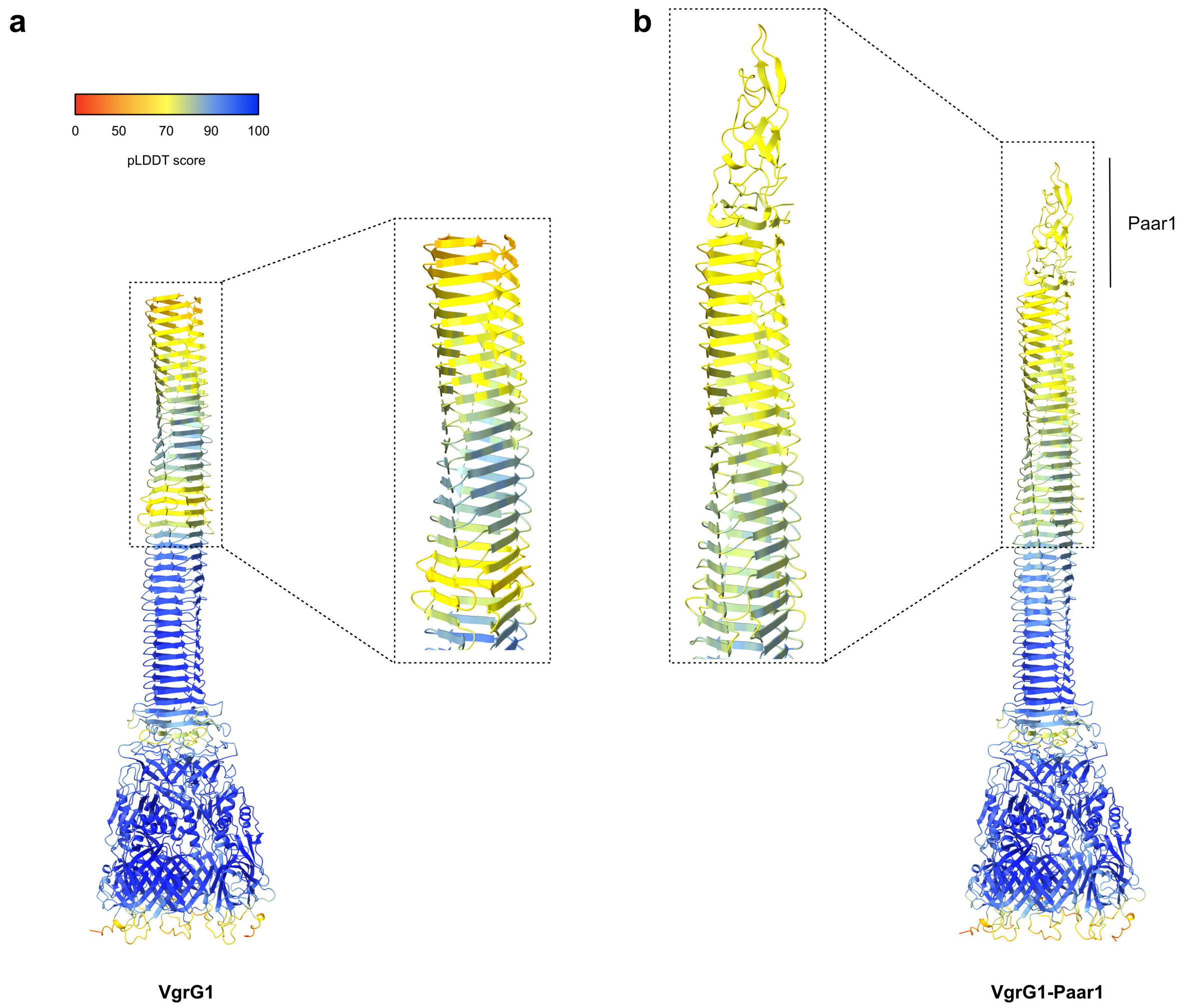


**Supplementary Figure 5**. **AlphaFold predictions of VgrG1 alone or in complex with Paar1.** Ribbon representation of predicted structures of VgrG1 from *Serratia marcescens* Db10, either **a)** alone (as in Figure 4), or **b)** in complex with Paar1. A zoom on the VgrG1 tip is provided to show how it could be stabilized by Paar1 protein. Predicted structures were generated by AlphaFold2 and coloured according to the pLDDT score, from red (pLDDT <50, lowest confidence) to blue (pLDDT >90, highest confidence).

#### Supplementary Figure 6

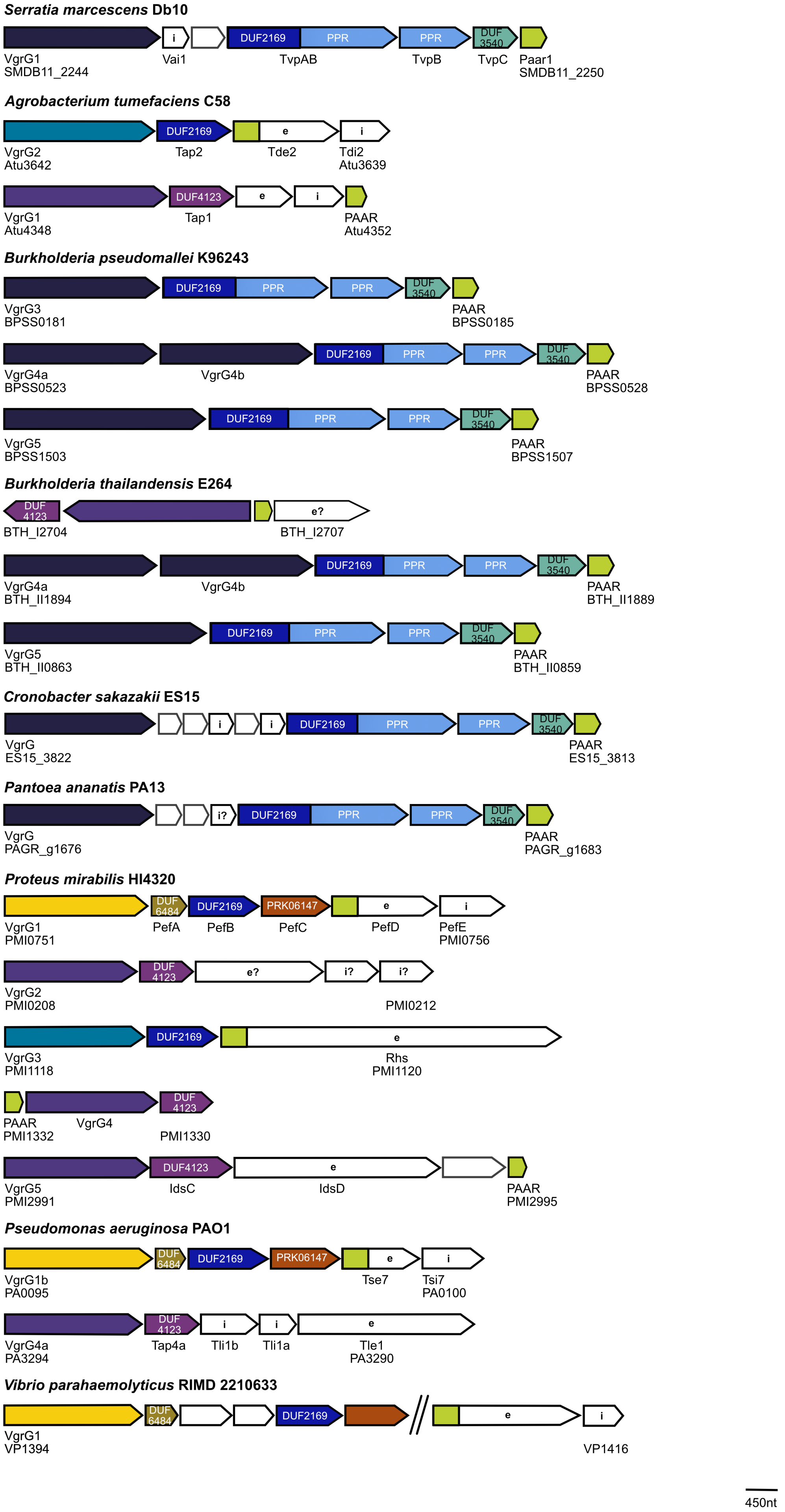

#### Supplementary Figure 6

**Supplementary Figure 6**. **Conservation of Tvp accessory proteins in representative Gram-negative species.** The genetic context of T6SS-associated DUF2169-containing accessory proteins in *Serratia marcescens* Db10 (accession: HG326223), *Agrobacterium tumefaciens* C58 (accession: AE007869.2), *Pantoea ananatis* PA13 (accession: CP003085.1), *Cronobacter sakazakii* ES15 (accession: AFK01427), *Proteus mirabilis* HI4320 (accession: NC_010554.1), *Pseudomonas aeruginosa* PAO1 (accession: AE004091.2), *Burkholderia pseudomallei* K96243 (accession: NC_006350.1 and NC_006351.1) and *B. thailandensis* E264 (accession: CP000085.1 and CP000086.1). If present in the genome, the genetic context of DUF4123-containing accessory proteins is also depicted. Each open reading frame (ORF) is represented by an arrow whose size is proportional to its gene length. The name and/or genomic identifier of the ORFs at each end of the set of genes is indicated below each set of genes. When identified, conserved domains (PPR for pentapeptide repeat, DUF3540, DUF6484, PRK06147) are highlighted inside the ORFs as well as effector (e) and immunity (i) encoding genes.

#### Supplementary Figure 7

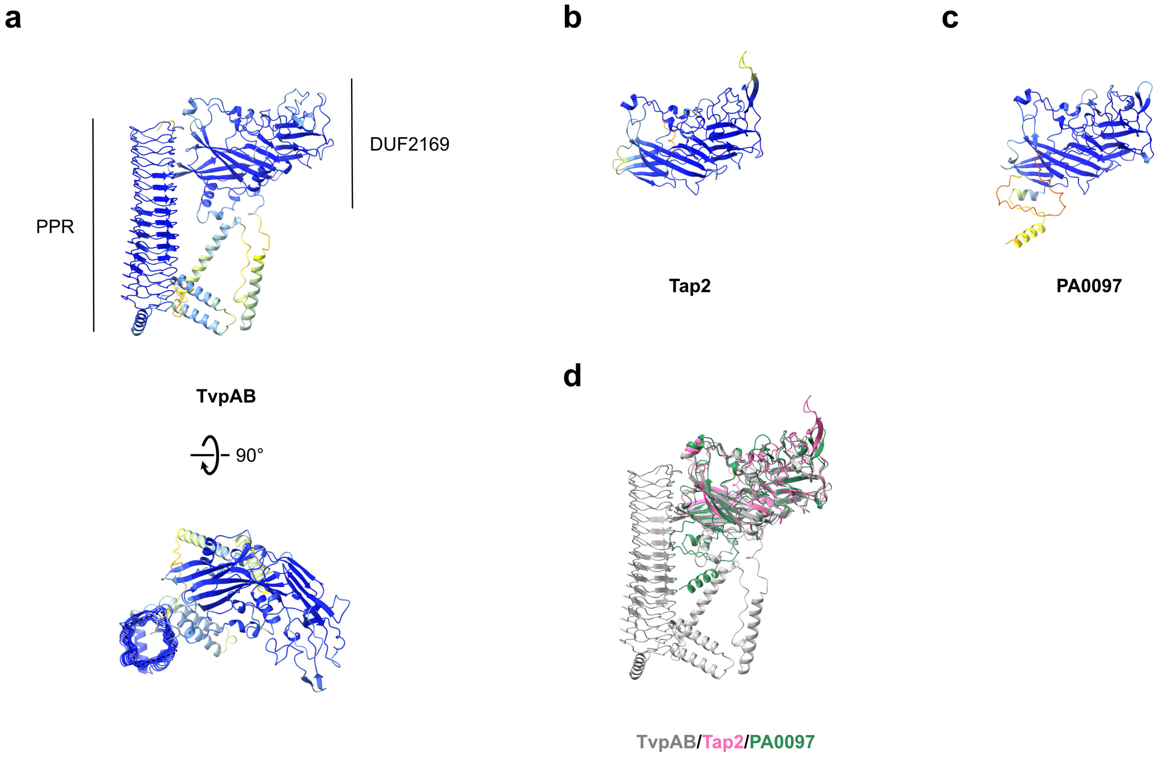


**Supplementary Figure 7. Predicted structures of TvpAB, Tap2 and PA0097 proteins. a)** Ribbon representation of the predicted structure of TvpAB from *Serratia marcescens* Db10. The DUF2169 and pentapeptide repeat (PPR) domains are highlighted. A 90° rotated view is provided to show the quadrilateral shape of the PPR β-helix. **b)** Ribbon representation of the predicted structure of the *Agrobacterium tumefaciens* DUF2169-containing accessory protein Tap2. **c)** Ribbon representation of the predicted structure of the *Pseudomonas aeruginosa* DUF2169-containing accessory protein PA0097. **d)** Superposition of TvpAB (grey), Tap2 (pink) and PA0097 (green) predicted structures. **a-c)** Predicted structures were generated by AlphaFold2 and coloured according to the pLDDT score, from red (pLDDT <50, lowest confidence) to blue (pLDDT >90, highest confidence).

### Supplementary Figure 8

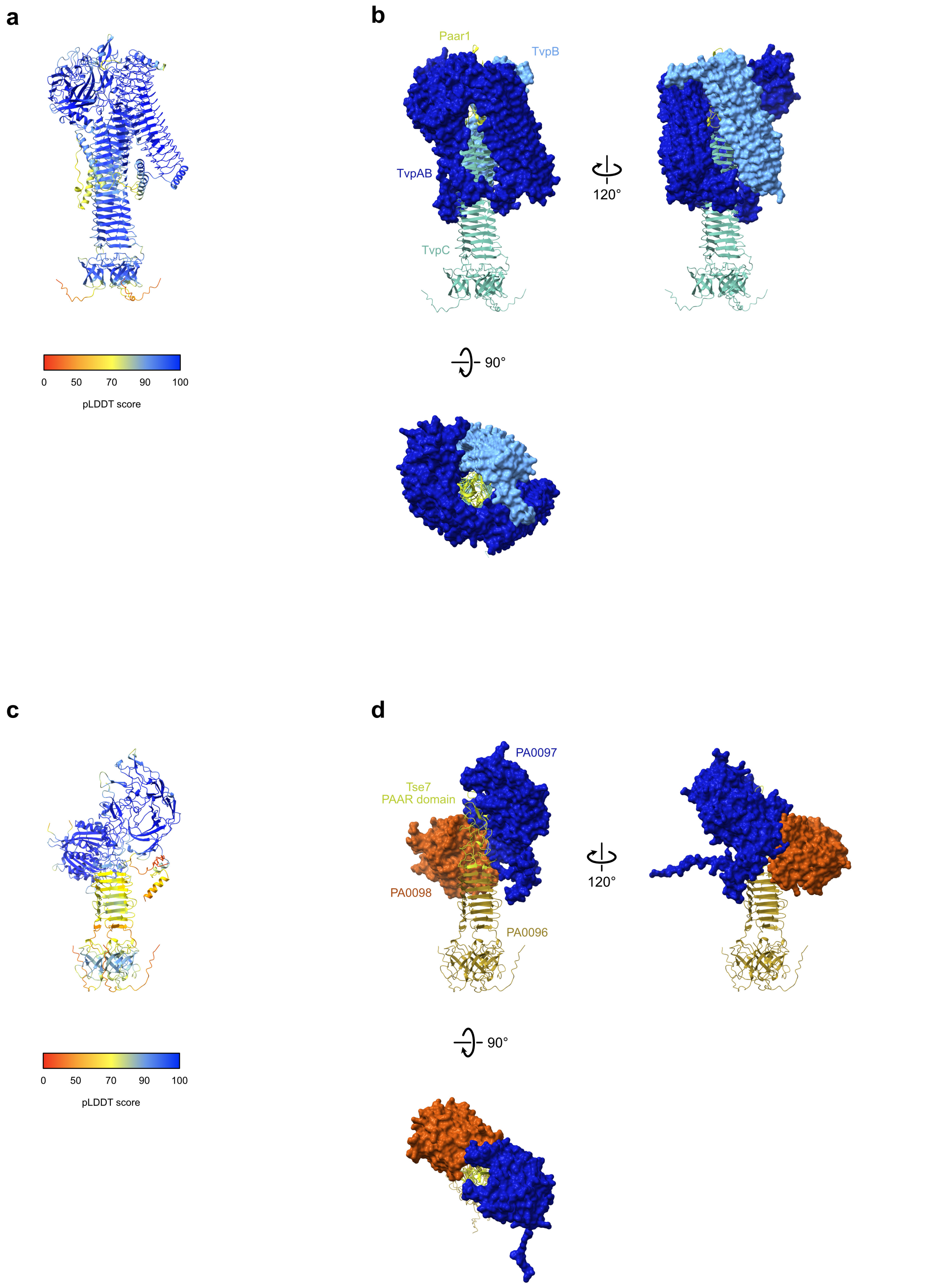


**Supplementary Figure 8**. **Predicted structures of *Serratia marcescens* and *Pseudomonas aeruginosa* accessory pre-complexes. a)** Ribbon representation of the *S. marcescens* Tvp accessory pre-complex coloured according to the pLDDT score from AlphaFold2 modelling, from red (pLDDT <50, lowest confidence)

to blue (pLDDT >90, highest confidence). **b)** The *S. marcescens* Tvp accessory pre-complex, as in Figure 6a, with the surface of TvpAB and TvpB highlighted to show how the proteins wrap the tip of the pre-complex. **c)** Ribbon representation of the *P. aeruginosa* accessory pre-complex coloured according to the pLDDT score from AlphaFold2 modelling, from red (pLDDT <50, lowest confidence) to blue (pLDDT >90, highest confidence). **d)** The *P. aeruginosa* accessory pre-complex, as in Figure 6b, with the surface of PA0097 and PA0098 highlighted to show how the proteins wrap the tip of the pre-complex.

#### Supplementary Figure 9

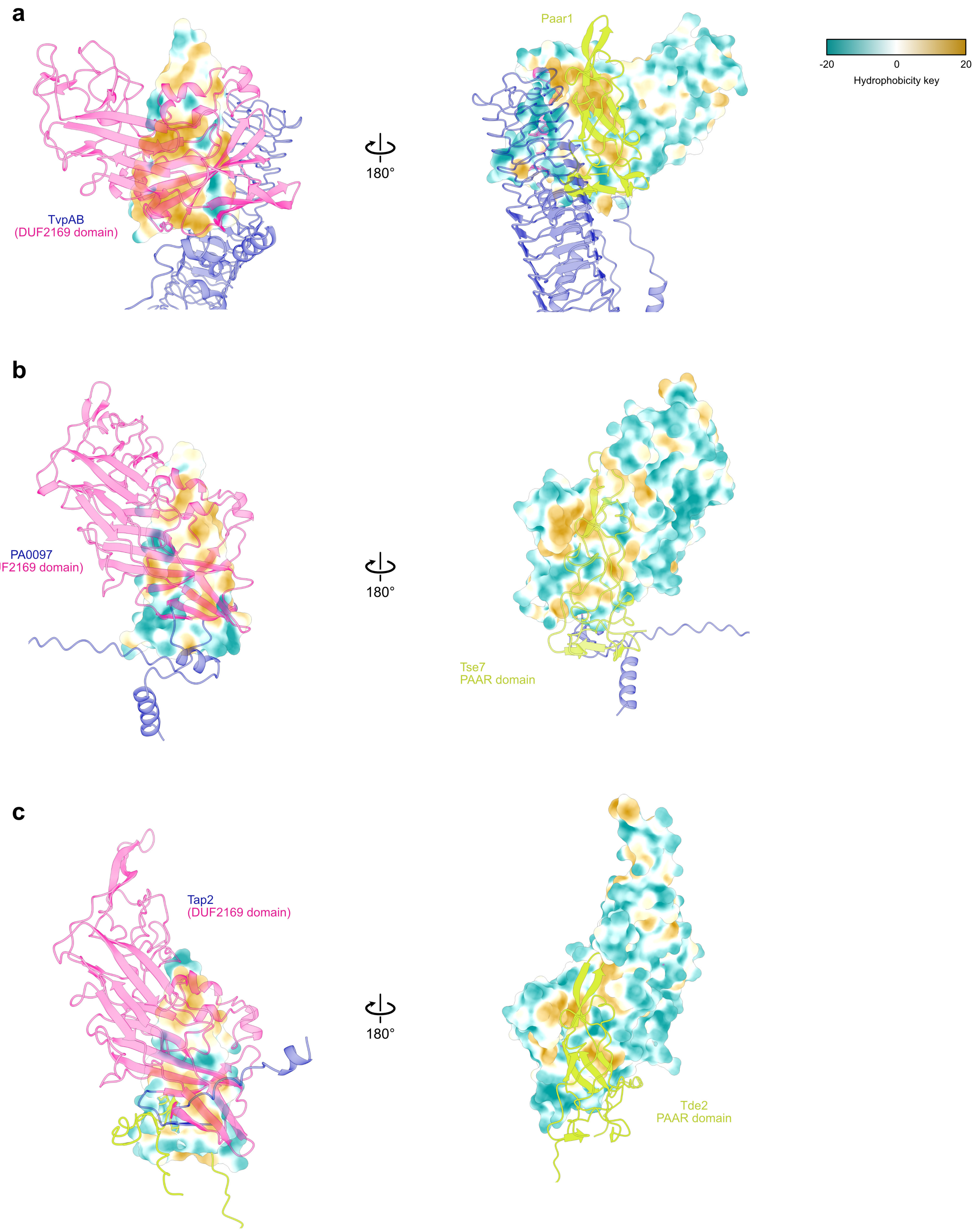


**Supplementary Figure 9. The interaction between the DUF2169 domain of TvpA/TvpAB proteins and the PAAR domain involves hydrophobic patches on both proteins. a)** Interaction between TvpAB and Paar1 from *Serratia marcescens* Db10. **b)** Interaction between PA0097 and the PAAR domain of Tse7 from *Pseudomonas aeruginosa* PA01. **c)** Interaction between Tap2 and the PAAR domain of Tde2 from

#### Supplementary Figure 9

*Agrobacterium tumefaciens* C58. **a-c)** Left: The hydrophobic surface of PAAR domains is highlighted and TvpA/TvpAB proteins are represented as ribbon with their DUF2169 domain colored in deep pink. Right: The hydrophobic surface of the DUF2169 domains of TvpA/TvpAB is highlighted whereas the PAAR domains are represented as ribbon and colored in light green. Hydrophobic surfaces are colored from dark cyan (most hydrophilic) to white to dark golden (most lipophilic) as in the key.

#### Supplementary Figure 10

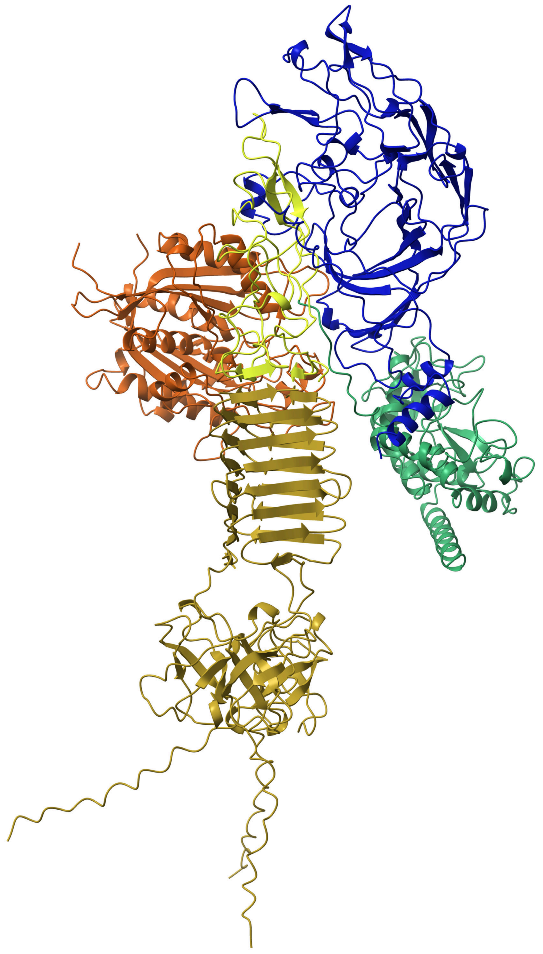

**Supplementary Figure 10. Predicted structure of the *Pseudomonas aeruginosa* accessory pre-complex with the full Tse7 effector.** Ribbon representation of the predicted structure of the *P. aeruginosa* accessory pre-complex generated with AlphaFold2. PA0096 is colored in ginger, PA0097 in medium blue, PA0098 in rust, the PAAR domain of Tse7 effector in light green and the effector domain of Tse7 in sea green.

#### Supplementary Figure 11

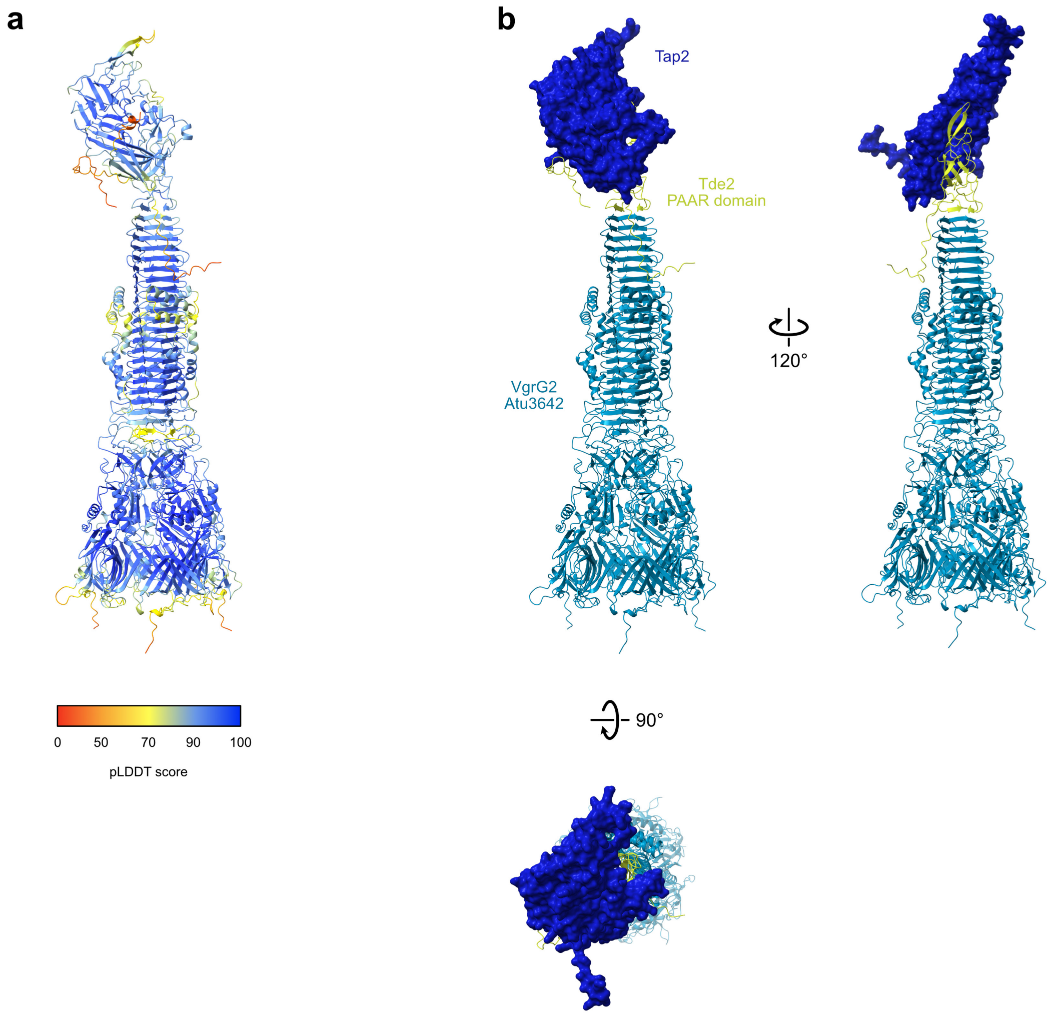


**Supplementary Figure 11. Predicted structure of the *Agrobacterium tumefaciens* VgrG2-associated accessory complex. a)** Ribbon representation of the *A. tumefaciens* VgrG2-associated accessory complex coloured according to the pLDDT score from AlphaFold2 modelling, from red (pLDDT <50, lowest confidence) to blue (pLDDT >90, highest confidence). **b)** The *A. tumefaciens* VgrG2-associated accessory complex, as in Figure 6e, with the surface of Tap2 highlighted to show how the protein wraps the tip of the complex.

### Supplementary Figure 12

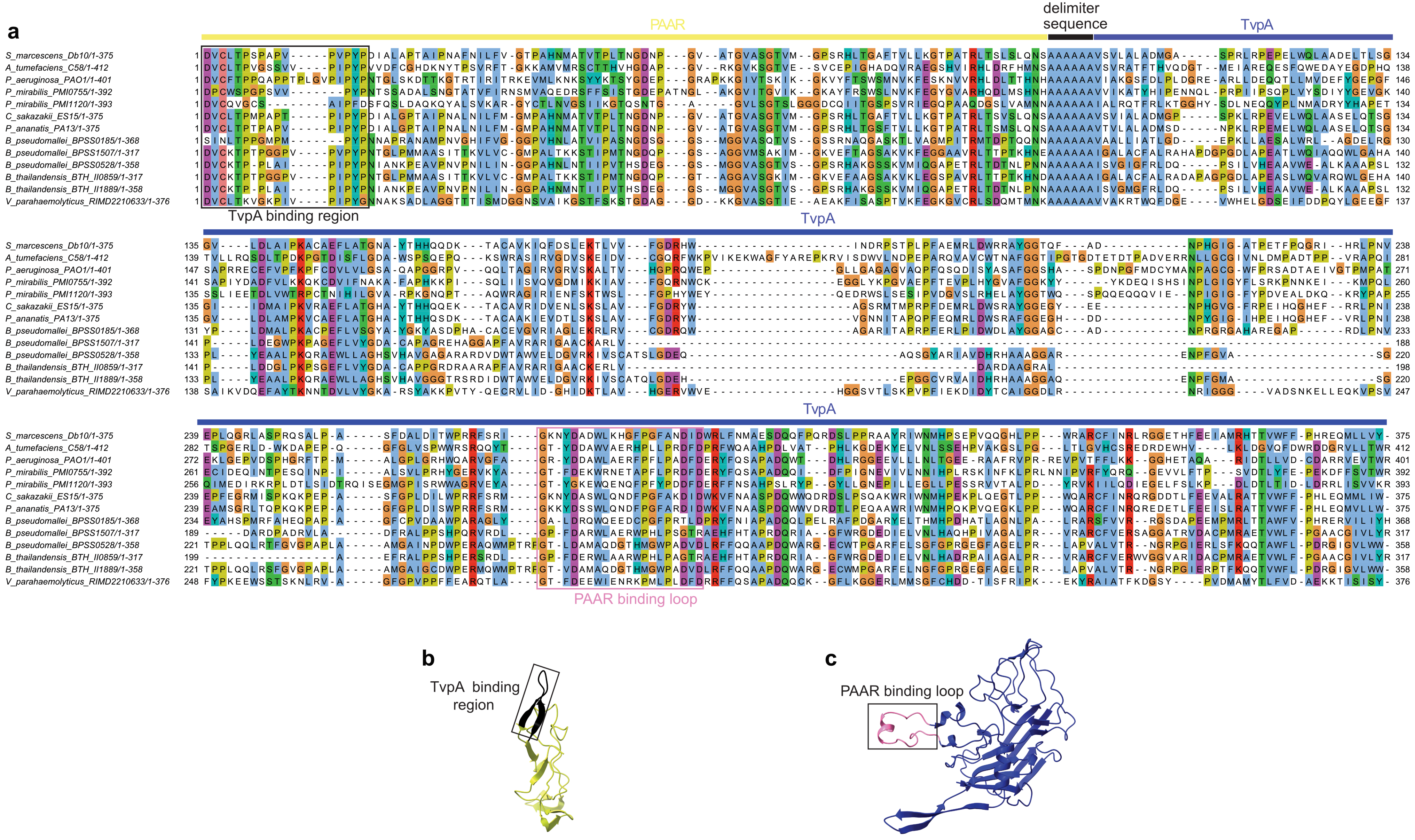

#### Supplementary Figure 12

**Supplementary Figure 12: Identification of conserved regions of the TvpA and PAAR DUF4150 proteins for phylogenetic analysis of the Tvp system. a)** A concatenated PAAR-TvpA sequence was used to generate a multiple sequence alignment using the MUSCLE algorithm[1] and coloured using ClustalX[2]. The alignment was also used for the purpose of phylogenetic tree construction. The chosen sequences were validated against structural comparisons between the corresponding AlphaFold3 predicted structures and each represents a conserved structural region. The conserved region is represented by: **b)** PAAR amino acids 16-110 (shown in green-yellow) and **c)** TvpA amino acids 26-299 (shown in medium-blue), both from *Serratia marcescens* Db10. Highlighted within these regions is the conserved TvpA binding interface of PAAR (black box in panel **a**, black ribbon in panel **b**) and the corresponding PAAR binding loop from TvpA (pink box in panel **a**, pink ribbon in panel **b**). Ribbon representations of structures in **b** and **c** were generated in ChimeraX[3].

#### Supplementary Figure 13

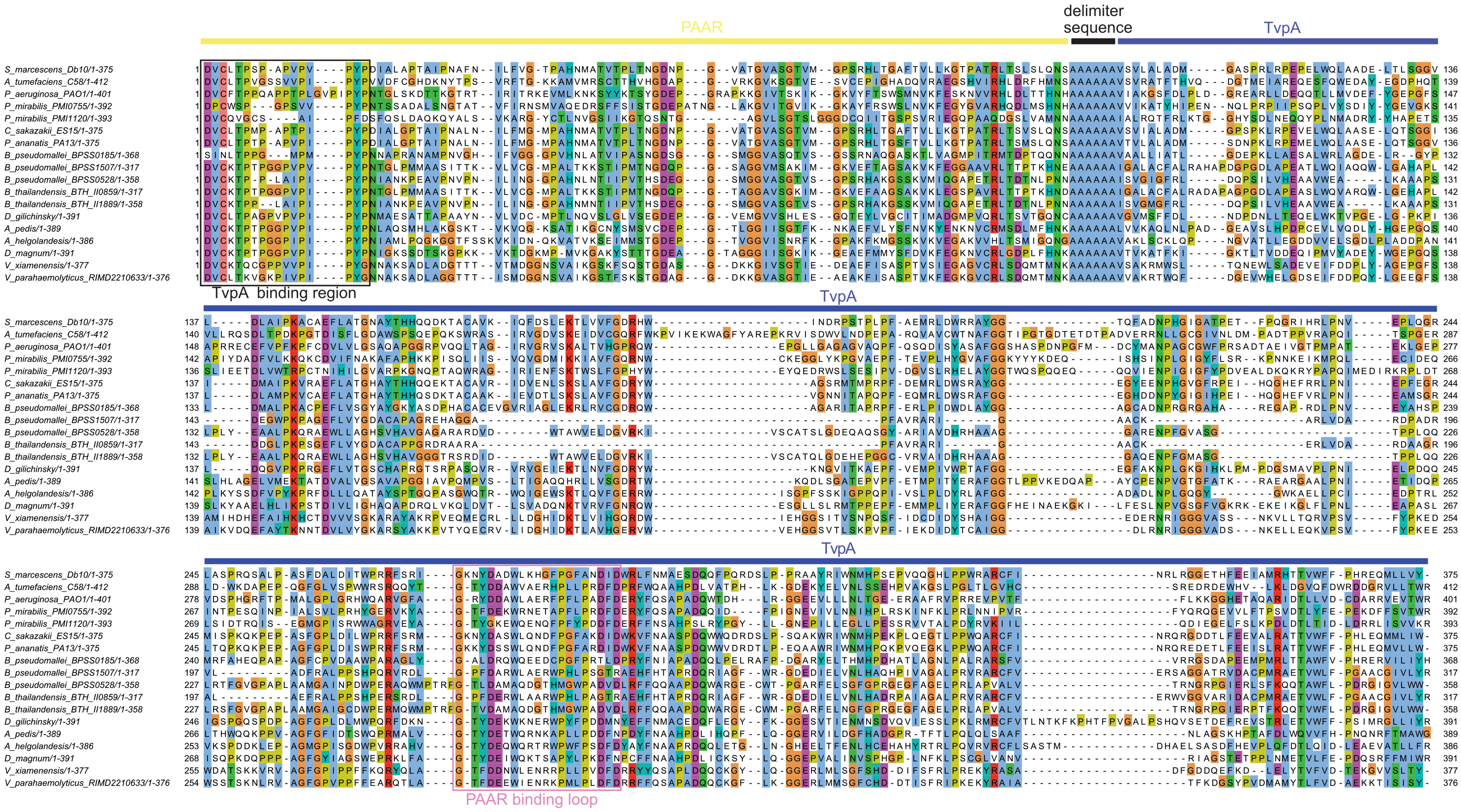

**Supplementary Figure 13: Alignment of an extended selection of PAAR, TvpA sequences for phylogenetic analysis.** As for Supplementary Figure 12, except that an expanded set of concatenated PAAR-TvpA sequences was selected to include bacterial species from additional phyla.

#### Supplementary Figure 14

a

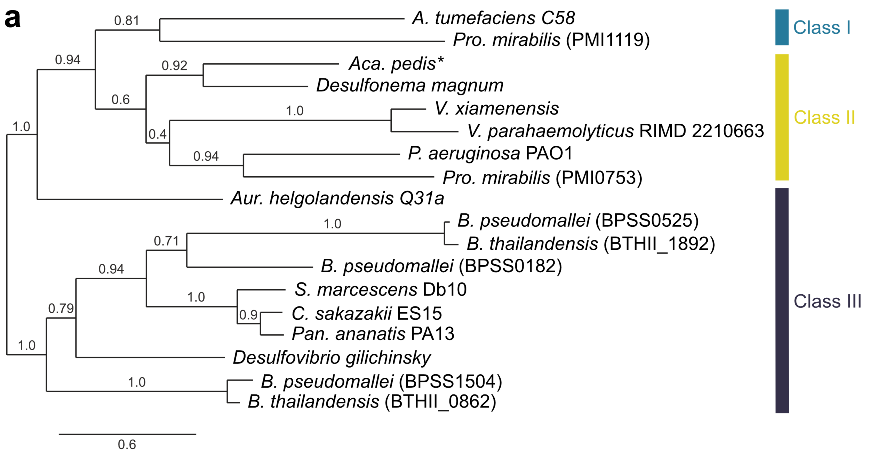


b

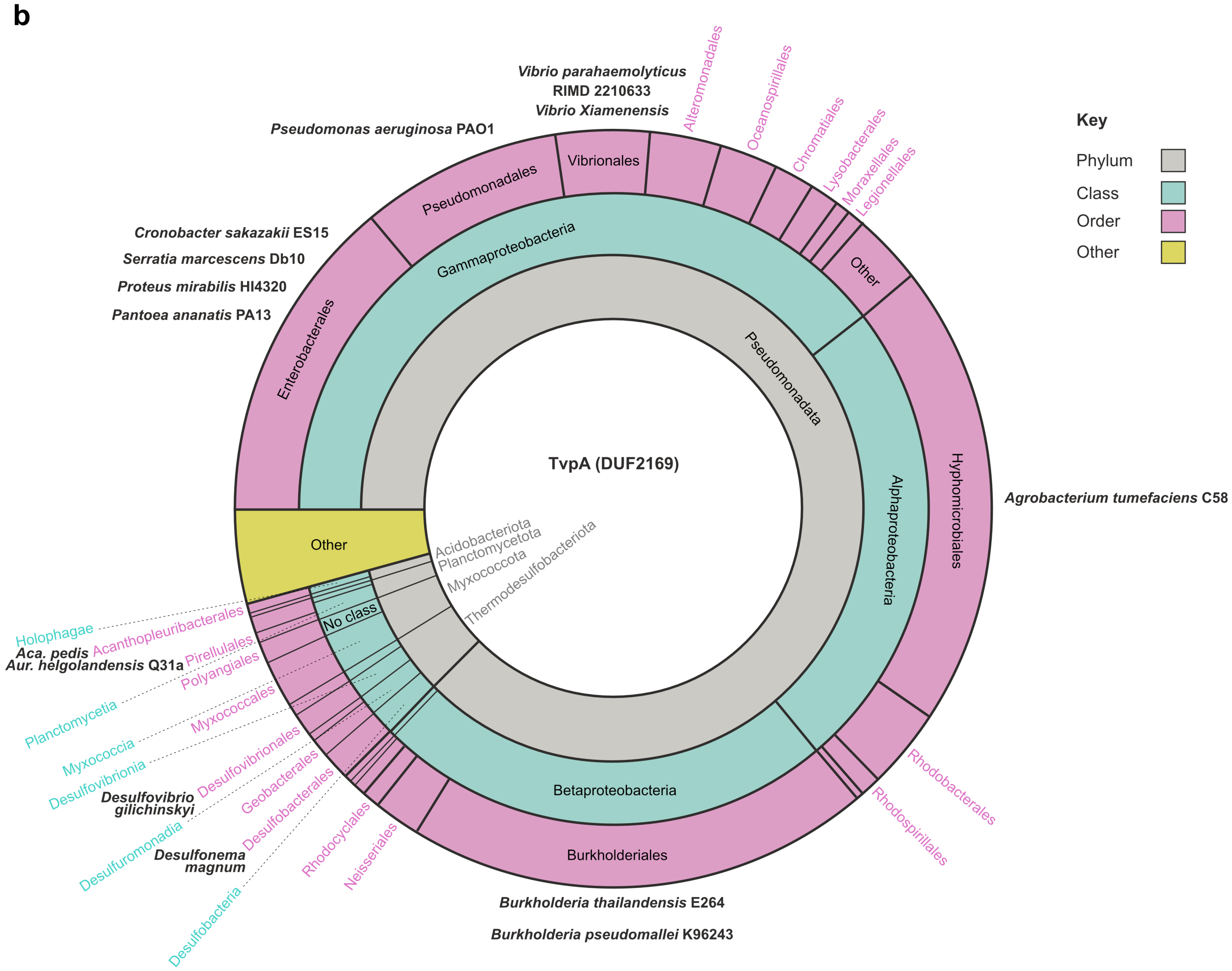


**Supplementary Figure 14. Extended phylogeny and taxonomy**. **a)** An extended phylogenetic tree, similar to that in Figure 7a, was constructed from the expanded set of sequences in the extended sequence alignment in Supplementary Figure 13. This represents an extension from the phylum

**Supplementary Figure 14**

*Pseudomonodata* to include representatives from additional phyla[4]. Where strains contain multiple Tvp systems, the accession number for the TvpA protein is enclosed in brackets to distinguish between them. *No syntenous TvpC protein. **b)** Taxonomic distribution of TvpA (DUF2169) genes (InterPro entry: IPR018683; Pfam: PF09937)[5,6] as in Figure 7d with the positions of *Acanthopleuribacter pedis* (Acidobacteriota), *Aureliella helgolandensis* (Planctomycetota), *Desulfonema magnum* and *Desulfovibrio gilichinskyi* (Thermodesulfobacteriota) indicated.

#### Supplementary Figure 15

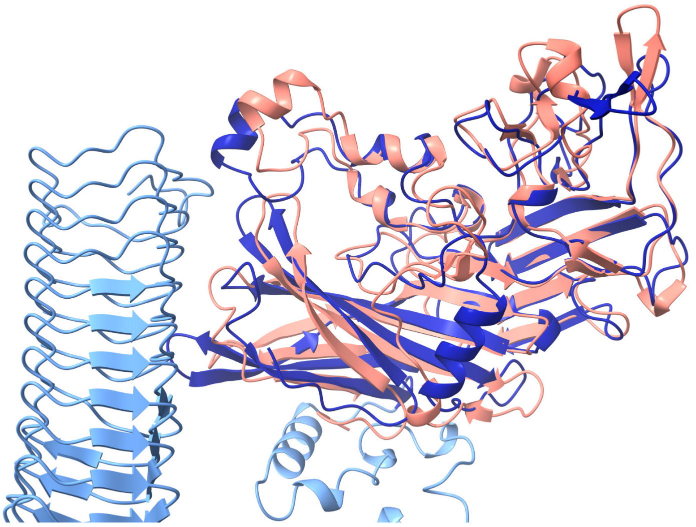

**Supplementary Figure 15**: **Comparison of the Alphafold3 predicted structure of TvpAB from *Serratia marcescens* Db10 with the experimentally-determined structure of TvpA of *Vibrio xiamenensis*.** Ribbon representations of the AlphaFold3 model of TvpAB from *S. marcescens* and the crystal structure of *V. xiamenensis* TvpA (PDB: 8VTH). Superimposition of the structures was performed using ChimeraX[3] (RMSD = 1.076 Å between 140 pruned atom pairs) and the figure focuses on the DUF2169 domain. The structure of the TvpA domain of Db10 (amino acids 1-323) is shown in dark blue while the rest of the structure is shown in light blue. The *V. xiamenensis* TvpA protein is shown in salmon.

#### Supplementary Figure 16

a c

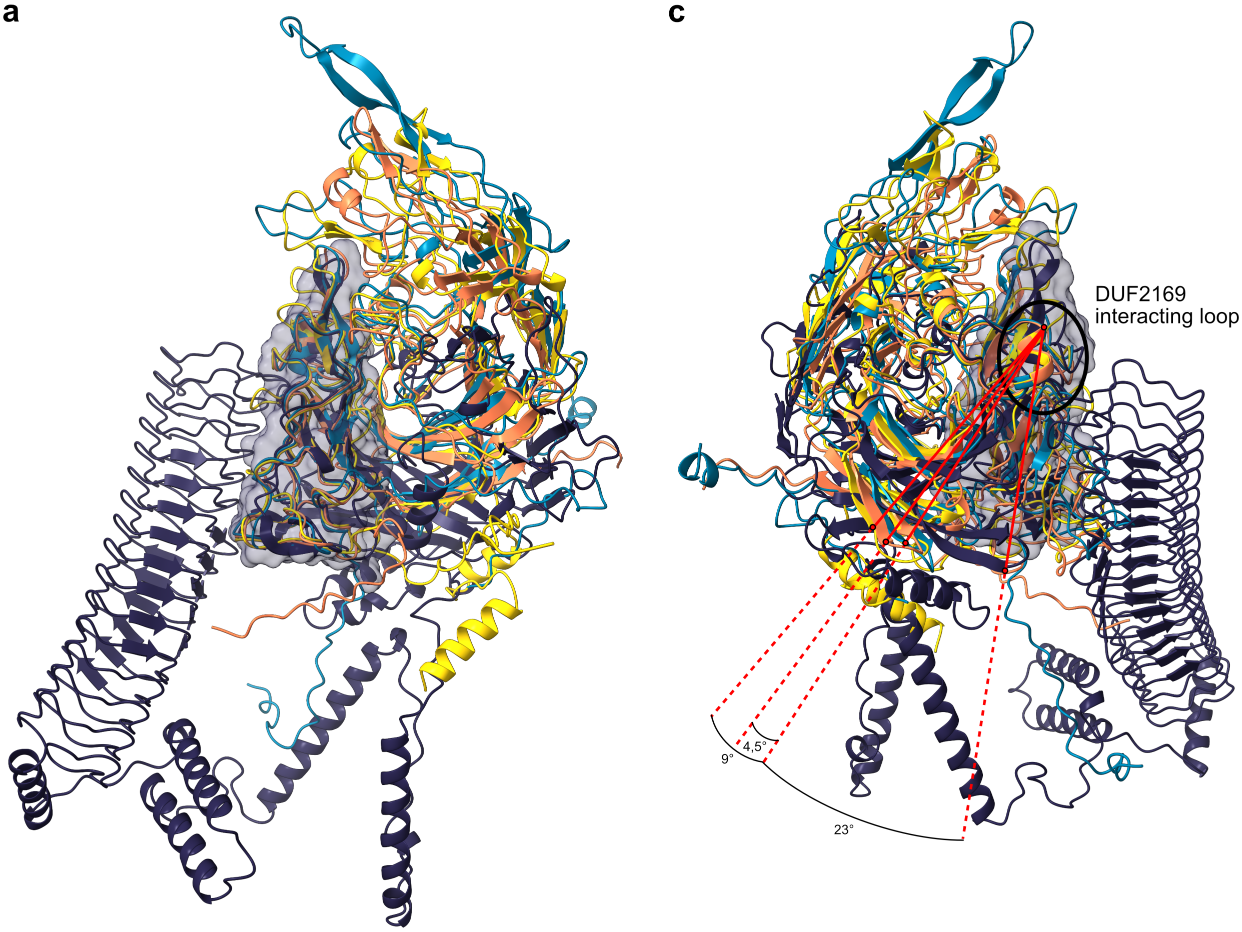


b

| Orientation compared to class I *A. tumefaciens* TvpA-PAAR complex | | Rotation | Shift |
|---|---|---|---|
| Class II | *P. aeruginosa* PAO1 | 3.6° | 0.1 Å |
| | *V. parahaemolyticus* RMID 2210633 | 11.7° | 1.4 Å |
| Class III | *S. marcescens* Db10 | 43.6° | -0.2 Å |

**Supplementary Figure 16 : Comparison of the orientation of the DUF2169 domain in the TvpA-PAAR complex between the different classes of Tvp systems. a)** AlphaFold3 structural predictions of TvpAB-Paar1 from *Serratia marcescens* Db10 (dark blue), Tap2-PAAR domain of Tde2 from *Agrobacterium tumefaciens* C58 (cyan), PA0097-PAAR domain of Tse7 from *Pseudomonas aeruginosa* PAO1 (yellow), and VP1398-PAAR domain of VP1415 from *Vibrio parahaemolyticus* RIMD 2210633 (salmon) are represented as cartoons and superimposed by their PAAR domain. The surface of Db10 Paar1 is highlighted. **b)** Table summarizing the rotation and shift required to realign the DUF2169 domain of all the TvpA proteins to the DUF2169 domain of Tap2 from *A. tumefaciens* starting from the superimposition presented in panel **a**. **c)** Same structural predictions as in panel **a** but superimposed by the short α-helix located in the interacting loop defined by Sachar *et al.*[7]. An axis was drawn between the top of this short α-helix and the end of the same β-strand for all the structures (red line). Flexibility of the DUF2169 domain among the different classes is estimated by the rotation angle formed between the axis of *A. tumefaciens* (Class I) and those of Class II (*P. aeruginosa* and *V. parahaemolyticus*) or Class III (*S. marcescens*).

### Supplementary Figure 17

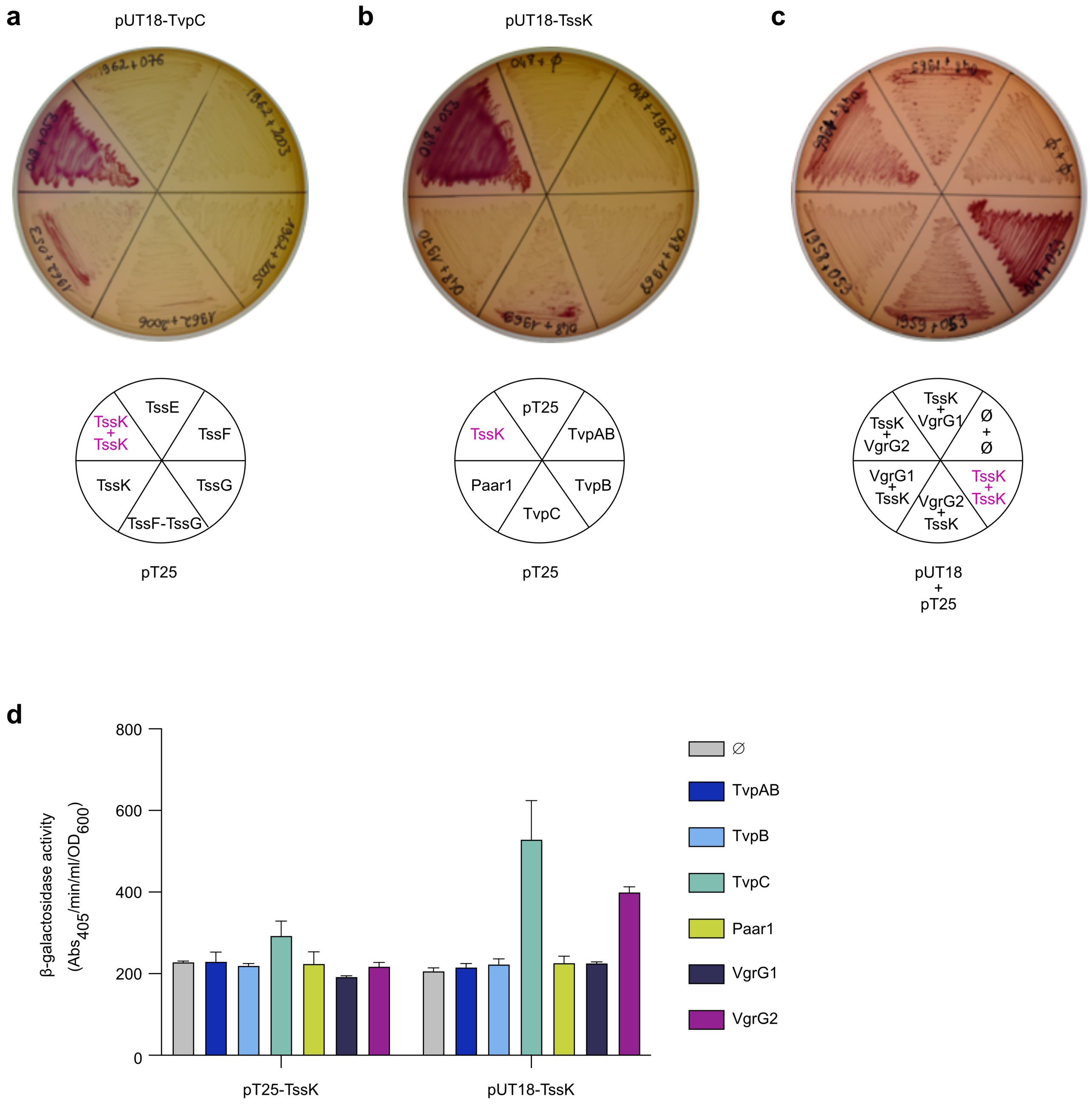


**Supplementary Figure 17**. **Interaction between TssK and TvpC by bacterial-two-hybrid assay.** **a-c)** Detection of interactions between T6SS components by observing growth of *E. coli* Δ*cya* carrying derivatives of the pUT18 and pT25 vectors directing the expression of T18 and T25 fusions with proteins of interest on MacConkey maltose plates. A red colour indicates positive interaction between proteins in the two-hybrid system. As a positive control, the interaction between TssK monomers was assayed, the negative control was empty pUT18 and pUT25 plasmids (Ø + Ø). **a)** Testing the interaction between TvpC (in pUT18) and baseplate components (in pT25) from *S. marcescens* Db10 when expressed in *E. coli* Δ*cya*. **b)** Testing the interaction between TssK (in pUT18) and Tvp accessory proteins (in pT25) from *S. marcescens* Db10 when expressed in *E. coli* Δ*cya*. **c)** Testing the interaction between TssK and VgrG proteins from *S. marcescens* Db10 (either in pUT18 or pT25) when expressed in *E. coli* Δ*cya*. **d)** Measurement of β-galactosidase activity to confirm and quantify the interactions between TssK and Tvp accessory proteins or VgrG proteins presented in panels **a**-**c**.

#### Supplementary Table 1

| Name | Description/Genotype | Source/reference |
|---|---|---|
| **Strains** | | |
| *Serratia marcescens* Db10 | | |
| Db10 | wild-type | [8] |
| SJC11 | Δ*tssE* (*SMDB11_2271*) | [9] |
| FRA01 | Δ*vgrG2* (*SMDB11_2276*) | [10] |
| FRA02 | Δ*vgrG1* (*SMDB11_2244*) | [10] |
| FRA03 | Δ*vgrG1* Δ*vgrG2* | [10] |
| DWL05 | Δ*vai1* (*SMDB11_2245)* | This study |
| DP10 | Δ*SMDB11_2246* | This study |
| DP09 | Δ*vai1* Δ*tssE* | This study |
| DP15 | Δ*vai1* Δ*tssE* Sm-resistant derivative | This study |
| DP13 | Δ*SMDB11_2246* Δ*tssE* | This study |
| DP16 | Δ*SMDB11_2246* Δ*tssE* Sm-resistant derivative | This study |
| DWL03 | Δ*vai1Δ2246* Δ*tssE* | This study |
| DWL08 | Δ*vai1-Δ2246* Δ*tssE* Sm-resistant derivative | This study |
| LM36 | Δ*vgrG2* Δ*vai1* | |
| LM32 | Δ*vgrG2* Δ*SMDB11*_2246 | This study |
| KK1 | Δ*tvpAB* (*SMDB11_2247*) | This study |
| DP11 | Δ*tvpB* (*SMDB11_2248*) | This study |
| DWL04 | Δ*tvpC* (*SMDB11_2249*) | This study |
| FRC32 | Δ*paar1* (*SMDB11_2250*) | [10] |
| DP02 | Δ*vgrG2* Δ*tvpAB* | This study |
| CE76 | Δ*vgrG2* Δ*tvpB* | This study |
| DP03 | Δ*vgrG2* Δ*tvpC* | This study |
| FRC26 | Δ*vgrG2* Δ*paar1* | [10] |
| LM33 | Δ*vgrG1* Δ*tvpAB* | This study |
| YL37 | Db10 Δ9 [Δ*ssp1* (Δ*SMDB11_2261*), Δ*ssp2* (Δ*SMDB11_2264*), Δ*ssp3/tfe1* (Δ*SMDB11_1112*), Δ*ssp4* (Δ*SMDB11_3980*), Δ*ssp5* (Δ*SMDB11_4628*), Δ*ssp6* (Δ*SMDB11_4673*), Δ*rhs1* (Δ*SMDB11_2278*), *rhs2*$_{H1369A}$ (*SMDB11_1610*$_{H1369A}$), Δ*slp* (Δ*SMDB11_0927*)] | This study; constituent mutations in [10–12] |
| GM131 | Δ9 Δ*vgrG1* | This study |
| CE030 | Δ9 Δ*vgrG2* | This study |
| CE036 | *vai1*-3xFlag (encodes *vai1* with a C-terminal 3xFLAG tag at native *vai1* locus in a wild-type background) | This study |
| CE043 | *vai1*-3xFlag Sm-resistant derivative | This study |
| CE049 | Δ*vgrG2* SMDB11_2245-3xFlag (encodes *vai1* with a C-terminal 3xFLAG tag at native *vai1* locus in a Δ*vgrG2* background) | This study |
| JAD09 | Δ*rhsI1* Δ*tssH* Sm-resistant derivative | [11] |
| AO01/JAD06 | Δ*sip4* Δ*ssp4* Sm-resistant derivative | [10] |
| LM26 | His-VgrG1 (encodes VgrG1 with an N-terminal $His_6$ tag in a wild-type background) | This study |
| LM66 | His-VgrG1 Δ*tvpAB* (encodes VgrG1 with an N-terminal $His_6$ tag in a Δ*tvpAB* background) | This study |
| CE56 | His-VgrG1 Δ*tvpC* (encodes VgrG1 with an N-terminal $His_6$ tag in a Δ*tvpC* background) | This study |
| CE31 | TvpAB-HA (encodes TvpAB with a C-terminal HA tag in a wild-type background) | This study |

### Supplementary Table 1

| | | |
|---|---|---|
| CE68 | TvpAB-HA Δ*tvpC* (encodes TvpAB with a C-terminal HA tag in a Δ*tvpC* background) | This study |
| CE59 | HA-TvpC (encodes TvpC with an N-terminal HA tag in a wild-type background) | This study |
| CE75 | HA-TvpC Δ*vgrG1* (encodes TvpC with an N-terminal HA tag in a Δ*vgrG1* background) | This study |
| CE70 | HA-TvpC Δ*tvpAB* (encodes TvpC with an N-terminal HA tag in a Δ*tvpAB* background) | This study |
| *Pseudomonas fluorescens* | | |
| KT02 | Sm-resistant derivative of *P. fluorescens 55* | [9] |
| *Serratia marcescens* SM39 | | |
| SM39 | Wild type, intrinsically Sm-resistant | [13] |
| *E. coli* | | |
| CC118λpir | Cloning host and donor strain for pKNG101-derived allelic exchange plasmids (λ*pir*) | [14] |
| MG1655 Δ*cyaA::apr* | Strain used for bacterial two hybrid assay | F. Sargent |

**Supplementary Table 1**. **List of bacterial strains used in this study.**

#### Supplementary Table 2

| Name | Description/Genotype | Source/reference |
|---|---|---|
| pKNG101 | Suicide vector for allelic exchange (Sm$^R$ *sacBR mobRK2 oriR6K*) | [15] |
| pSC3048 | Modified version of the recombineering, λ Red recombinase-encoding pORTMAGE plasmid. The dominant negative mutant *mutL* gene has been removed to prevent off-target effects due to suppression of DNA repair during integration of oligonucleotide CE75. The origin of replication from pBAD18-Kn was introduced in place of the conjugation and replication machinery to allow for the stable maintenance of the plasmid in *S. marcescens* Db10 (Kn$^R$) | This study |
| pSUPROM | Vector for constitutive expression of cloned genes under the control of the *E. coli tat* promoter (Kn$^R$) | [16] |
| pUT18 | Bacterial Two Hybrid plasmid (for fusion of protein of interest with C-terminal T18 fragment of CyaA; ApR) | [17] |
| pT25 | Bacterial Two Hybrid plasmid (for fusion of protein of interest with N-terminal T25 fragment of CyaA; CmR) | [18] |
| pSC1914 | pSUPROM *SMDB11_2245* | This study |
| pSC1915 | pSUPROM *SMDB11_2246* | This study |
| pSC1916 | pSUPROM *tvpAB* (*SMDB11_2247*) | This study |
| pSC1917 | pSUPROM *tvpB* (*SMDB11_2248*) | This study |
| pSC1918 | pSUPROM *tvpC* (*SMDB11_2249*) | This study |
| pSC1919 | pSUPROM *paar1* (*SMDB11_2250*) | This study |
| pSC601 | pKNG101-derived allelic exchange plasmid for the generation of chromosomal in-frame Δ*vgrG2* (Δ*SMDB11_2276*) deletion | [10] |
| pSC602 | pKNG101-derived allelic exchange plasmid for the generation of chromosomal in-frame Δ*vgrG1* (Δ*SMDB11_2244*) deletion | [10] |
| pSC921 | pKNG101-derived allelic exchange plasmid for the generation of chromosomal in-frame Δ*SMDB11_2245* deletion | This study |
| pSC937 | pKNG101-derived allelic exchange plasmid for the generation of chromosomal in-frame Δ*SMDB11_2246* deletion | This study |
| pSC577 | pKNG101-derived allelic exchange plasmid for the generation of chromosomal in-frame Δ*SMDB11_2245-2246* deletion | This study |
| pSC902 | pKNG101-derived allelic exchange plasmid for the generation of chromosomal in-frame Δ*tvpAB* deletion (Δ*SMDB11_2247*) | This study |
| pSC945 | pKNG101-derived allelic exchange plasmid for the generation of chromosomal in-frame Δ*tvpB* deletion (Δ*SMDB11_2248*) | This study |
| pSC923 | pKNG101-derived allelic exchange plasmid for the generation of chromosomal in-frame Δ*tvpC* deletion (Δ*SMDB11_2249*) | This study |
| pSC3031 | pKNG101-derived allelic exchange plasmid for chromosomal replacement of native vai1 with vai1with 3xFLAG (C-terminal tag) | This study |
| pSC2026 | pKNG101-derived allelic exchange plasmid for chromosomal replacement of native *vgrG1* gene with His-VgrG1 (N-terminal tag) | This study |
| pSC3027 | pKNG101-derived allelic exchange plasmid for chromosomal replacement of native *tvpAB* gene with TvpAB-HA (C-terminal tag) | This study |
| pSC3088 | pKNG101-derived allelic exchange plasmid for chromosomal replacement of native *tvpC* gene with HA-tvpC (N-terminal tag) | This study |
| pSC048 | pUT18 *tssK* (*SMDB11_2253*) | [10] |
| pSC1962 | pUT18 *tvpC* (*SMDB11_2249*) | This study |
| pSC1958 | pUT18 *vgrG1* (*SMDB11_2244*) | This study |
| pSC1959 | pUT18 *vgrG2* (*SMDB11_2276*) | This study |
| pSC053 | pT25 *tssK* (*SMDB11_2253*) | [10] |
| pSC076 | pT25 *tssE* (*SMDB11_2271*) | This study |

**Supplementary Table 2**

| | | |
|---|---|---|
| pSC1967 | pT25 *tvpAB* (*SMDB11_2247*) | This study |
| pSC1968 | pT25 *tvpB* (*SMDB11_2248*) | This study |
| pSC1969 | pT25 *tvpC* (*SMDB11_2249*) | This study |
| pSC1970 | pT25 *paar1* (*SMDB11_2250*) | This study |
| pSC1965 | pT25 *vgrG1* (*SMDB11_2244*) | This study |
| pSC1966 | pT25 *vgrG2* (*SMDB11_2276*) | This study |
| pSC2003 | pT25 *tssF* (*SMDB11_2272*) | This study |
| pSC2005 | pT25 *tssG* (*SMDB11_2273*) | This study |
| pSC2006 | pT25 *tssF-tssG* (*SMDB11_2272-73*) | This study |

**Supplementary Table 2. List of plasmids used in this study.**

#### Supplementary Table 3

| Plasmid | Sequence of relevant primers (5'-3')[a,b] | Description |
|---|---|---|
| pSC1914 | TATAGGATCCATGTCGAAGGCCAAAAATGTTTTCA | Forward primer to clone SMDB11_2245 into pSUPROM (BamHI) |
| | TATATCTAGATCAAAGCCGGGAAAGCTCTTCGATC | Reverse primer to clone SMDB11_2245 into pSUPROM (XbaI) |
| pSC1915 | TATAGGATCCATGGGCGCGTTCATTGCCTTTTCGA | Forward primer to clone SMDB11_2246 into pSUPROM (BamHI) |
| | TATATCTAGATCAAGAGGACAGTGCGTTGTCCTTC | Reverse primer to clone SMDB11_2246 into pSUPROM (XbaI) |
| pSC1916 | TATAGGATCCATGAAAGTCGTAAAACCTCTGCGCC | Forward primer to clone *tvpAB* (SMDB11_2247) into pSUPROM (BamHI) |
| | TATATCTAGATCATACTTCCTTCCCGTTGTGGCGA | Reverse primer to clone *tvpAB* (SMDB11_2247) in pSUPROM (XbaI) |
| pSC1917 | TATATCTAGAATGACCTCAAAACCAGAGCAGATAC | Forward primer to clone *tvpB* (SMDB11_2248) in pSUPROM (XbaI) |
| | TATAAAGCTTTCAAATTCGGCTCTGGCGCTGTGCG | Reverse primer to clone *tvpB* (SMDB11_2248) in pSUPROM (HindIII) |
| pSC1918 | TATAGGATCCATGAGCGTTGCCAACACAACTCGCG | Forward primer to clone *tvpC* (SMDB11_2249) in pSUPROM (BamHI) |
| | TATATCTAGATTATCCGACGTGGATTTGTTCTGAA | Reverse primer to clone *tvpC* (SMDB11_2249) in pSUPROM (XbaI) |
| pSC1919 | TATAGGATCCATGTTTGCCAACAGCCAAATGATCG | Forward primer to clone *paar1* (SMDB11_2250) in pSUPROM (BamHI) |
| | TATATCTAGATCAGGGCGCAAGCAGCAACACTTTT | Reverse primer to clone *paar1* (SMDB11_2250) in pSUPROM (XbaI) |
| pSC1962 | TATAAAGCTTGTAAAGAGGAGGAGTA**ATGAGCGTTGCCAACACAAC** | Forward primer to clone ***tvpC*** (SMDB11_2249) in pUT18 (HindIII) |
| | TATATCTAGAGA**TCCGACGTGGATTTGTTCTG** | Reverse primer to clone ***tvpC*** (SMDB11_2249) without stop in pUT18 (XbaI) |
| pSC1958 | TATACTGCAGGTAAAGAGGAGGAGTA**ATGGATCGTCTTATCATTGC** | Forward primer to clone ***vgrG1*** (SMDB11_2244) in pUT18 (PstI) |
| | TATATCTAGAGA**CAAGAAGATCGCCAGCGCCT** | Reverse primer to clone ***vgrG1*** (SMDB11_2244) without stop in pUT18 (XbaI) |
| pSC1959 | TATAAAGCTTGTAAAGAGGAGGAGTA**ATGTTGGACCGCATTATTGC** | Forward primer to clone ***vgrG2*** (SMDB11_2276) in pUT18 (HindIII) |
| | TATATCTAGAGA**GGATTTCGGGAACAGCTGAT** | Reverse primer to clone ***vgrG2*** (SMDB11_2276) without stop in pUT18 (XbaI) |
| pSC076 | TATAGGATCCA**ATGAACGATAAACGCCCACC** | Forward primer to clone ***tssE*** (SMDB11_2271) in pT25 (BamHI) |
| | TATAGGTACC**TTACCCTATGTCTTTCAGATCGAAGTG** | Reverse primer to clone ***tssE*** (SMDB11_2271) in pT25 (KpnI) |
| pSC1967 | TATAGGATCCG**AAAGTCGTAAAACCTCTGCG** | Forward primer to clone ***tvpAB*** (SMDB11_2247) without start in pT25 (BamHI) |
| | TATACCCGGG**TCATACTTCCTTCCCGTTGT** | Reverse primer to clone ***tvpAB*** (SMDB11_2247) in pT25 (SmaI) |
| pSC1968 | TATACTGCAGGA**ACCTCAAAACCAGAGCAGAT** | Forward primer to clone ***tvpB*** (SMDB11_2248) without start in pT25 (PstI) |
| | TATAGGTACC**TCAAATTCGGCTCTGGCGCT** | Reverse primer to clone ***tvpB*** (SMDB11_2248) in pT25 (KpnI) |
| pSC1969 | TATAGGATCCG**AGCGTTGCCAACACAACTCG** | Forward primer to clone ***tvpC*** (SMDB11_2249) without start in pT25 (BamHI) |
| | TATACCCGGG**TTATCCGACGTGGATTTGTT** | Reverse primer to clone ***tvpC*** (SMDB11_2249) in pT25 (SmaI) |
| pSC1970 | TATAGGATCCG**TTTGCCAACAGCCAAATGAT** | Forward primer to clone ***paar1*** (SMDB11_2250) without start in pT25 (BamHI) |

#### Supplementary Table 3

| | | |
|---|---|---|
| | TATACCCGGG**TCAGGGCGCAAGCAGCAACA** | Reverse primer to clone ***paar1*** (SMDB11_2250) in pT25 (SmaI) |
| pSC1965 | TATACTGCAGGA**GATCGTCTTATCATTGCGCA** | Forward primer to clone ***vgrG1*** (SMDB11_2244) without start in pT25 (PstI) |
| | TATAGGTACC**TTACAAGAAGATCGCCAGCG** | Reverse primer to clone ***vgrG1*** (SMDB11_2244) in pT25 (KpnI) |
| pSC1966 | TATAAGATCTG**TTGGACCGCATTATTGCGC** | Forward primer to clone ***vgrG2*** (SMDB11_2276) without start in pT25 (BglII) |
| | TATAGATATC**TCAGGATTTCGGGAACAGCT** | Reverse primer to clone ***vgrG2*** (SMDB11_2276) in pT25 (EcoRV) |
| pSC2003 | TATAGGATCCG**GACAGTAAACTGTTAGATTA** | Forward primer to clone ***tssF*** (SMDB11_2272) without start in pT25 (BamHI) |
| | TATACCCGGG**TCATATCAGCGTCCTTTTGC** | Reverse primer to clone ***tssF*** (SMDB11_2272) in pT25 (SmaI) |
| pSC2005 | TATAGGATCCG**AACGCGGAGATGATCGCCAC** | Forward primer to clone ***tssG*** (SMDB11_2273) without start in pT25 (BamHI) |
| | TATACCCGGG**TCAGAACGGGTTTTCCAGCG** | Reverse primer to clone ***tssG*** (SMDB11_2273) in pT25 (SmaI) |
| pSC2006 | TATAGGATCCG**GACAGTAAACTGTTAGATTA** | Forward primer to clone ***tssF-tssG*** (SMDB11_2272-73) without start for *tssF* in pT25 (BamHI) |
| | TATACCCGGG**TCAGAACGGGTTTTCCAGCG** | Reverse primer to clone ***tssF-tssG*** (SMDB11_2272-73) in pT25 (SmaI) |
| pSC921 | TATATCTAGAATCAGTCTTTCAAGCTTGACGG | Forward primer to clone upstream region of SMDB11_2245 for allelic exchange (XbaI) |
| | TATAACTAGTCGACATGGGAGTCCCTTTACAAG | Reverse primer to clone upstream region of SMDB11_2245 for allelic exchange (SpeI) |
| | TATAACTAGTGAGCTTTCCCGGCTTTGAG | Forward primer to clone downstream region of SMDB11_2245 for allelic exchange (SpeI) |
| | TATAGGGCCCACAGTGCGTTGTCCTTCCC | Reverse primer to clone downstream region of SMDB11_2245 for allelic exchange (ApaI) |
| pSC937 | TATATCTAGACCAATTTCAAGCTCAGTAATGATCC | Forward primer to clone upstream region of SMDB11_2246 for allelic exchange (XbaI) |
| | TATAACTAGTGCCCATCGTGACACTTCC | Reverse primer to clone upstream region of SMDB11_2246 for allelic exchange (SpeI) |
| | TATAACTAGTAACGCACTGTCCTCTTGATCATCC | Forward primer to clone downstream region of SMDB11_2246 for allelic exchange (SpeI) |
| | TATAGGGCCCGGATTATCGGCGAACTGG | Reverse primer to clone downstream region of SMDB11_2246 for allelic exchange (ApaI) |
| pSC577 | TATATCTAGAATCAGTCTTTCAAGCTTGACGG | Forward primer to clone upstream region of SMDB11_2245 for allelic exchange (XbaI) |
| | TATAACTAGTCGACATGGGAGTCCCTTTACAAG | Reverse primer to clone upstream region of SMDB11_2245 for allelic exchange (SpeI) |
| | TATAACTAGTAACGCACTGTCCTCTTGATCATCC | Forward primer to clone downstream region of SMDB11_2246 for allelic exchange (SpeI) |
| | TATAGGGCCCGGATTATCGGCGAACTGG | Reverse primer to clone downstream region of SMDB11_2246 for allelic exchange (ApaI) |
| pSC902 | TGTATCTAGACTGTTTACCGTCTTTGGCATCC | Forward primer to clone SMDB11_2247 upstream region for allelic exchange (XbaI) |
| | TGTAGGATCCGACTTTCATTGTGTGTGTAAAGCTTCC | Reverse primer to clone SMDB11_2247 upstream region for allelic exchange (BamHI) |
| | TATAGGATCCACATTGCCTCGCCACAACG | Forward primer to clone SMDB11_2247 downstream region for allelic exchange (BamHI) |
| | TATAGGGCCCAATTGGTTATCGGTAAACTGGC | Reverse primer to clone SMDB11_2247 downstream region for allelic exchange (ApaI) |

#### Supplementary Table 3

| | | |
|---|---|---|
| pSC945 | TATATCTAGACACGCTGGCCAACATTACC | Forward primer to clone upstream region of SMDB11_2248 for allelic exchange (XbaI) |
| | TATAACTAGTTGAGGTCATACTTCCTTCCCG | Reverse primer to clone upstream region of SMDB11_2248 for allelic exchange (SpeI) |
| | TATAACTAGTAGCCGAATTTGAATGCATTGAC | Forward primer to clone downstream region of SMDB11_2248 for allelic exchange (SpeI) |
| | TATAGGGCCCGTAATCTTCCGCCTGATAATCC | Reverse primer to clone downstream region of SMA2248 for allelic exchange (ApaI) |
| pSC923 | TGTATCTAGAAGTTTGAAGGCGTGAACTTCG | Forward primer to clone upstream region of SMDB11_2249 for allelic exchange (XbaI) |
| | TATAACTAGTAACGCTCATGGTGATCCCCTC | Reverse primer to clone upstream region of SMDB11_2249 for allelic exchange (SpeI) |
| | TATAACTAGTATCCACGTCGGATAACCTGC | Forward primer to clone downstream region of SMB11_2249 for allelic exchange (SpeI) |
| | TATAGGGCCCGTTGACTTCACTGCTGATG | Reverse primer to clone downstream region of SMDB11_2249 for allelic exchange (ApaI) |
| pSC2026 | TATAAAGCTT**CATCATCATCATCATCAC**GATCGTCTTATCATTGCGCAC | Fwd primer to incorporate Nter **His Tag** on VgrG1 (SMDB11_2244) by allelic exchange (HindIII) |
| | TATAGTCGACGTCAGAAACGCTCCAGCTATTG | Reverse primer within VgrG1 coding region to incorporate Nter His Tag on VgrG1 (SMDB11_2244) by allelic exchange (SalI) |
| pSC3088 | gggcccggtacccagctt | Forward primer for pBluescript to integrate HA-SMDB11_2249 encoding sequence at the *ApaI* site |
| | tctagagcggccgccacc | Reverse primer for pBluescript to integrate HA-SMDB11_2249 encoding sequence at the *XbaI* site |
| | gcggtggcggccgctctagaCAGTTTGAAGGCGTGAAC | Forward primer for upstream region of SMDB11_2249. Overlap with pBluescript shown as lower case. |
| | **TGCATAATCAGGAACATCATAAGGATA**CATGGTGATCCCCTCGTG | Reverse primer for upstream region of SMDB11_2249. HA tag encoding sequence shown in bold. |
| | **TATCCTTATGATGTTCCTGATTATGCA**AGCGTTGCCAACACAACTC | Forward primer for coding region of SMDB11_2249. HA tag encoding sequence shown in bold. |
| | aaaagctgggtaccgggcccGGTGATGTCTTTTGAGGTAATGAG | Reverse primer for coding region of SMDB11_2249. Overlap with pBluescript shown as lower case. |
| Primer CE75 | T*T*CGAAACCGTTGGTCAAACGCACACGGCAAACTTTACGCAGTGCGGAGTTTGGTTTGTTCGGGGTAGTGGTATATACGCGGGTGCATA*C*G | $Sm^R$ SNP from DB11 to introduce into target strains. Stars denote phosphothiorate bonds to increase stability and resistance to nucleases |
| **Plasmid** | **Details of synthetic insert** | |
| pSC3031 | Synthetic insert of DNA sequence encoding a C-terminal 3xFLAG tag (shown in **bold**) on Vai1 (SMDB11_2245) to be introduced into the native genomic locus by allelic exchange (containing *XbaI*/*ApaI* sites (underlined) for sub-cloning into pKNG101). Produced by GeneArt (ThermoFisher):<br>TCTAGACACACGATGCTGGATATCTCGCATAACCCGCTGTCGATCAGCAAAACCAATTTCAAGCTCAGTAATGATCCCATGACGTTCACTCAGCTTGGCGCGCTGGTCACCAACGGTGCCTTACGATTGAGCACCAAGGCGCTGGCGATCTTCTTGTAAAGGGACTCCCATGTCGAAGGCCAAAAATGTTTTCACCGTTATCGGGTTGATTTTATTGGCTCCCATCCTCAGTCTGGTGGGTTTTTTCAAATTCATCACGGTCGATATCTGGGGTTATTTCAAACATATCGGTTCTTCCCTCAGCGTGATGGAGAACGGTATTTCCGCTTCCGGGCAAATCGTCAGCATTCGCCAGACCAACCTGTGGGACGGAAATCGGCCGGTCTGCGAAGTCGAAGTCAAGTACATCGCCCAAGACGGAAAAAACCATACGGCCGTGGCGAAAGGCCCGATCAGCGTCGTGGATCTGCCGCGTTATCAGCCGGGCCATTTCACCGCGATAAAGTACGATCCGAAGAACCCCAAAAAAGCGGTGATCGAAGAGCTTTCCCGGCTT**GACTACAAAGACCATGACGGTGATTATAAAGATCATGATATCGATTACAAGGATGACGATGACAAG**TGAGCGAAAAGGATCCCTTCAATTACGCGATCCGATTTCAACGTTCTGCAAACGGCTTATCGTTCATCAGTTCGCT | |

| | |
|---|---|
| | AAATCACCTTGCATAAGGAAGTGTCACGATGGGCGCGTTCATTGCCTTTTCGACGCAGGACAGCCTCTGGGGACAGA<br>TCCTGTTTACCGTCTTTGGCATCCTGGTCGGCTGCGTATTGCTCGCTGCGATCGGCAAAACCTTTTTGTCGGCAAGCCG<br>AGCGGCCGCCTTACCCATGCGGTTGCTGATCTTTGTCGCGCTGATGCTGGCGCTCCTGAAATACAATCAGCAGGTGAT<br>GTCTGCGGTGAATGCTCCCGCAGAGCTGATCGGGTTCCTGATTTTGCCCGGTTTTTTGATGCGTTACCTCGGTGCGGC<br>GACACCCGGCAGACTGACGCTGGCGATAGAGTTGCCCTGTTTTGCGGCGCTCGTCGCGCTGCTGGTTCTGAGCACCG<br>CCGGCGTCGGATTTTTTGCGCCCTATCACCTGGAGAGCACCGGCCAATTTACCGCCAAGGCGATCGCCGCGTCATTGT<br>CCGTCGCGGGCCC |
| pSC3027 | Synthetic insert of DNA sequence encoding a C-terminal HA tag (shown in **bold**) on TvpAB (SMDB11_2247) to be introduced into the native genomic locus by allelic exchange (containing *XbaI*/*ApaI* sites (underlined) for sub-cloning into pKNG101). Produced by GeneArt (ThermoFisher):<br><br>TCTAGAGACATTCAGCCACGCCCGGCTCGACAAAGTCAGCTTCGTCAAATGCCGGTTGCTGGCGGTGAACTTCAGTC<br>AGGCGCGGCTGGAAAGTTGCGCCTGGGTTGACACCGAGACGCAAAGCCTCAGCTTCCACGCCGCACGTCTCACGGC<br>CTGCGCCTTTGCAGCGAAGACACTGTTGCCGCAGGCGGATTTCAGCGACGCGACGCTAAACCAGTGCAATTTACGCC<br>AGATGCCGTTGCAGCGGGCCAATTTTAGCCGGGCGCGTCTCAATAATTGCGATCTGTCGGAAACCCGATTGAACGAG<br>GCCGATTTCCGCCAGGCCAACGGCAGTGGCAGCCTGTTCATCCGCAGCGATTTGTCTCGCGCCAGCCTGCGCGATGC<br>CAATTTCATTGCGGCTATCCTGCAAAAATGCGTGCTGTCAGGCGCCGACCTGCAGGGCGCCAATCTGTTCCGGGCGG<br>ACCTGTCGCAGTCGCAAGTGGATCAGGCAACCCGGCTCGAGAGCGCCTATACCGCCAGGGTGAAAACATTGCCTCGC<br>CACAACGGGAAGGAAGTA**TATCCTTATGATGTTCCTGATTATGCA**TGACAACGGGAAGGAAGTATGACCTCAAAACC<br>AGAGCAGATACGCCAGCGCGTAAAACGCGGCGAGCCGATCGCCGGAGAGGACTTGCACGGCCTGTCATTCGCCGGT<br>CTGGATTTGGCCGGCGGCATGTTCAACGAACTGAACCTGAACGGCGTGAACTTCTCTGACTGCGATCTGCGCGACAG<br>CGTTTTCAGCGATTGCCAGCTCGAACACGCACAATTTACGCGGGCGGATCTGAAGCAAACGGCCTTCAATCAGTGCG<br>CCATGTCGGCCAGCCGCTTCAGCGAGAGCCATATCGAACTGACGATGTTCAACGATTGCCGGCTTGAACAGAGCGAC<br>TTCAGCCGGCTGTCGCTCAACCAAAGTCACTGGATGTCGTGCCAATTGGCCGGCGCGAATTTTTCCGCCACTCAGCAC<br>GATCGCACTACCTTTTACGAAAGCCCGCTCGACGGCGCCATGTTGAATCAGGCGCGCTTATGCCTTGTCACGTTCTTTC<br>GATTAAATCTCTGTAAAACGCAGTTTGAAGGCGTGAACTTCGACCGCGTCACGTTTTGGGCCC |

[a] Incorporated restriction sites for cloning into the respective vector are underlined.
[b] Sequences encoding protein epitope/affinity tags are in bold.

**Supplementary Table 3. Oligonucleotide primers and synthetic gene fragments used for plasmid construction.**

#### Supplementary Table 4

| Protein/Complex | pLDDT | pTM | ipTM |
|---|---|---|---|
| ***Serratia marcescens* Db10** | | | |
| VgrG1 | 88 | n.a. | n.a. |
| VgrG2 | 91.3 | n.a. | n.a. |
| TvpC | 93.1 | n.a. | n.a. |
| TvpAB | 92.8 | n.a. | n.a. |
| Db10 VgrG1-PAAR | 86.5 | 0.792 | 0.776 |
| Db10 pre-complex (TvpAB-TvpB-TvpC-PAAR) | 92.3 | 0.898 | 0.903 |
| ***Agrobacterium tumefaciens* C58** | | | |
| VgrG2 (Atu3642) | 92.1 | n.a. | n.a. |
| Tap2 | 93.1 | n.a. | n.a. |
| VgrG2 full complex (VgrG2-Tap2-PAAR Tde2) | 87.9 | 0.796 | 0.778 |
| ***Pseudomonas aeruginosa* PAO1** | | | |
| VgrG1b (PA0095) | 93.4 | n.a. | n.a. |
| PA0096 | 77 | n.a. | n.a. |
| PA0097 | 89.8 | n.a. | n.a. |
| PAO1 pre-complex (PA0096-PA0097-PA0098-PAAR Tse7) | 84.3 | 0.789 | 0.753 |

**Supplementary Table 4. Summary of the confidence scores of the Alphafold predictions presented in Figures 4 and 5.** Overall confidence scores of each AlphaFold prediction (pLDDT) and specific confidence scores for complexes (pTM and ipTM) are indicated. The PAAR domain of Tde2 corresponds to amino acids 1-172, and the PAAR domain of Tse7 corresponds to amino acids 1-140.

#### Supplementary Table 5

| Protein | β prism length (Å) | pLDDT |
|---|---|---|
| ***Serratia marcescens* Db10** | | |
| VgrG1 | 186 | 88 |
| VgrG2 | 81 | 91.3 |
| ***Agrobacterium tumefaciens* C58** | | |
| VgrG2 (Atu3642) | 96 | 92.1 |
| VgrG1 (Atu4348) | 118 | 84.9 |
| ***Pseudomonas aeruginosa* PAO1** | | |
| VgrG1b (PA0095) | 154 | 93.4 |
| VgrG1a (PA0091) | 76 | 89.6 |
| VgrG1c (PA2685) | 117 | 86.1 |
| VgrG2a (PA1511) | 110 | 83.7 |
| VgrG2b (PA0262) | 116 | 80.1 |
| VgrG3 (PA2373) | 117 | 88.2 |
| VgrG4a (PA3294) | 80 | 82.1 |
| VgrG4b (PA3486) | 80 | 76.8 |
| VgrG5 (PA5090) | 80 | 78.9 |
| VgrG6 (PA5266) | 80 | 82.5 |
| ***Burkholderia pseudomallei* K96243** | | |
| VgrG3 (BPSS0181) | 95 | 80.2 |
| VgrG4a (BPSS0523) | X | 71.6 |
| VgrG4b (BPSS0524) | X | 71.5 |
| VgrG5 (BPSS1503) | 165 | 65.6 |
| VgrG (BPSS2056) | 148 | 86.1 |
| VgrG (BPSS0958) | 89 | 86.9 |
| VgrG2 (BPSS0105) | 95 | 73 |
| VgrG6 (BPSS2093) | 152 | 87.6 |
| ***Burkholderia thailandensis* E264** | | |
| VgrG4a (BTH_II1894) | X | 72.6 |
| VgrG4b (BTH_II1893) | X | 70.3 |
| VgrG5 (BTH_II0863) | 163 | 66.3 |
| VgrG (BTH_II1436) | 89 | 86.9 |
| VgrG (BTH_I2705) | 105 | 75.6 |
| VgrG2 (BTH_II0129) | 95 | 75.6 |
| VgrG6 (BTH_II0265) | 151 | 87.8 |
| VgrG (BTH_II1531) | 105 | 77.1 |
| VgrG (BTH_I2693) | 109 | 71.9 |
| VgrG (BTH_I2697) | 105 | 76.4 |
| ***Pantoea ananatis* PA13** | | |
| VgrG (PAGR_g1676) | 110 | 80.6 |
| VgrG (PAGR_g1684) | 75 | 83.6 |
| ***Cronobacter sakazakii* ES15** | | |
| VgrG (ES15_3822) | 159 | 82.1 |
| VgrG (ES15_3808) | 76 | 87.1 |
| VgrG (ES15_3826) | 99 | 86 |
| VgrG (ES15_2015) | 109 | 86.5 |
| VgrG (ES15_2809) | 100 | 76.4 |
| ***Proteus mirabilis* HI4320** | | |
| VgrG1 (PMI0751) | 85 | 80.3 |

#### Supplementary Table 5

| | | |
|---|---|---|
| VgrG2 (PMI0208) | 80 | 83.5 |
| VgrG3 (PMI1118) | 76 | 81.2 |
| VgrG4 (PMI1331) | 80 | 83.8 |
| VgrG5 (PMI2991) | 99 | 82.5 |
| ***Vibrio parahaemolyticus* RIMD 2210633** | | |
| VgrG1 (VP1394) | 125 | 88.9 |
| VgrG (VPA1026) | n.d. | n.d. |

**Supplementary Table 5. Summary Table of VgrG β-prism length measurements.** Structural models of all the VgrG proteins in the set of eight representative bacteria (*S. marcescens* Db10, *A. tumefaciens* C58, *P. ananatis* PA13, *C. sakazakii* ES15, *P. mirabilis* HI4320, *P. aeruginosa* PAO1, *B. pseudomallei* K96243 and *B. thailandensis* E264) and in *V. parahamolyticus* RIMD 2210633 were generated using AlphaFold3. Each model was visualized with PyMol to measure the length of the β-prism when possible. These measurements were used to generate Figure 7c. Confidence scores (pLDDT) for each AlphaFold prediction are also indicated. n.d : not determined.
